# Purinergic receptor P2RY14 cAMP signaling regulates EGFR-driven Schwann cell precursor self-renewal and nerve tumor initiation in neurofibromatosis

**DOI:** 10.1101/2021.09.24.461701

**Authors:** Jennifer Patritti-Cram, Jianqiang Wu, Shinji Kuninaka, Robert A. Coover, Robert F. Hennigan, Tilat A. Rizvi, Katherine E. Chaney, Ramya Ravindran, Jose A. Cancelas, Robert J. Spinner, Nancy Ratner

**Affiliations:** Division of Experimental Hematology and Cancer Biology, Cancer & Blood Diseases Institute, Cincinnati Children’s Hospital Medical Center, Cincinnati, OH 45229, USA; Neuroscience Graduate Program, University of Cincinnati College of Medicine, Cincinnati, OH 45267-0713, USA; Division of Gene Regulation, Institute for Advanced Medical Research, Keio University, Tokyo 108-8345, Japan; Molecular and Developmental Biology, Cincinnati Children’s Hospital, Cincinnati, OH 45229, USA; Department of Neurosurgery, Mayo Clinic, Rochester, Minnesota 55902, USA; Department of Pediatrics, University of Cincinnati College of Medicine, Cincinnati, OH, 45267, USA; Hoxworth Blood Center, College of Medicine, University of Cincinnati, Cincinnati, OH 45229, USA; Dept. of Basic Pharmaceutical Sciences, High Point University, High Point, NC 27268, USA

**Author notes:** Corresponding Author: Nancy Ratner, Cincinnati Children’s Hospital Medical Center, 3333 Burnet Avenue, Cincinnati, OH 45229. **Lead Contact**: Further information and requests for resources and reagents should be directed to and will be fulfilled by the Lead Contact, Nancy Ratner, PhD.

**Keywords:** Neurofibromatosis type 1, NF1, Schwann cell-precursors, Schwann cell, P2RY14, neurofibromas, self-renewal, cAMP, EGFR, p-Akt, p-PKA peripheral nerve, tumorigenesis

## Abstract

Neurofibromatosis type 1 (NF1) is a genetic disorder characterized by nerve tumors called neurofibromas, in which Schwann cells (SCs) lack NF1 and show deregulated RAS signaling. NF1 is also implicated in regulation of cAMP. Gene expression profiling and protein expression identified P2RY14 in SCs and SC precursors (SCPs) implicating P2RY14 as a candidate upstream regulator of cAMP in EGF-dependent SCP. We found that SCP self-renewal was reduced by genetic or pharmacological inhibition of P2RY14. In *NF1* deficient SCs and malignant peripheral nerve sheath tumor (MPNST) cells, P2RY14 inhibition decreased EGFR-driven phospho-Akt and increased cAMP signaling. In a neurofibroma mouse model, genetic deletion of P2RY14 increased mouse survival, delayed neurofibroma initiation and rescued cAMP signaling. Conversely, elevation of cAMP diminished SCP number *in vitro* and diminished SC proliferation in neurofibroma bearing mice *in vivo*. These studies identify the purinergic receptor P2RY14 as a critical G-protein-coupled receptor (GPCR) in *NF1* mutant SCPs and SCs and suggest roles for EGFR-GPCR crosstalk in facilitating SCP self-renewal and neurofibroma initiation via cAMP and EGFR-driven phospho-Akt.

## Introduction

Neurofibromatosis type 1 (NF1) is an autosomal dominant disease that affects up to 1:2,000 individuals worldwide (Kallionpää et al., 2018). To date, there is no cure for NF1, which is characterized by multiple, variable, clinical manifestations (Friedman, 1998; Tabata et al., 2020). At least half of the children with NF1 develop plexiform neurofibromas (PNs), which are tumors within peripheral nerves. PN may be present at birth and show most rapid growth during the first decade of a child’s life (Nguyen, et al., 2012). PNs can occur in any cranial or peripheral nerve and have the potential to transform into lethal malignant peripheral nerve sheath tumors (MPNST) (Prudner, Ball, Rathore, & Hirbe, 2020). Neurofibroma infiltration of normal nerves in NF1 patients results in a complicated risk profile because it can cause nerve damage and compress nearby vital organs (Kim, et al., 2017). Therefore, understanding how neurofibromas form and how to treat them is under intense investigation.

Peripheral nerve glial cells, Schwann cells (SCs), are the only cell type in neurofibromas that show bi-allelic loss-of-function mutations in the *NF1* tumor suppressor gene (Serra et al., 1997, 2001). Neurofibroma SCs also show aberrant properties *ex vivo*, consistent with it being the primary pathogenic cell type in neurofibromas (Sheela et al., 1990; Kim et al., 1995). In mice, neural crest cells develop into Schwann cell precursors (SCPs) between embryonic day 11 (E11) and E13 (Jessen & Mirsky, 2019). SCPs or related boundary cap cells can serve as cells-of-origin for neurofibromas, as loss of *Nf1* in these cells causes plexiform neurofibroma formation (Zhu et al., 2002; Wu et al., 2008; Chen, et al., 2014; Chen et. al., 2019).

*In vitro*, embryonic SCPs retain multi-lineage differentiation potential and self-renewal capabilities for several passages, indicating that they are progenitor-like cells (Jessen & Mirsky, 2019). Mouse SCPs also express epidermal growth factor receptor^+^ (EGFR^+^) and respond to EGF with limited self-renewal (Williams et al. 2008). EGFR^+^ cells that co-express the SC marker S100 account for about 1.8% human neurofibroma cells (DeClue, et al., 2000). The idea that these cells may be tumor-initiating cells is consistent with the finding that human neurofibromas sorted for co-expression of the SC marker p75^+^ and EGFR show limited self-renewal *in vitro*. Also, EGFR-dependent *Nf1^-/-^* SCPs show increased self-renewal and form neurofibromas upon transplantation (Joseph et al., 2008; Williams et al. 2008). Together, these studies suggest the presence of progenitor-like cells in neurofibromas, which depend on EGFR for self-renewal.

EGFR signaling may also play additional roles in neurofibroma SCs. SCPs differentiate into SCs which when associated with a single large-diameter axonal segment become myelinating SCs. SCs associated with multiple smaller diameter axons become non-myelinating Remak cells (Mirsky et al., 2008). Neurofibroma SCs show a dramatic change in Remak Schwann cell morphology, bundling 1 or 2 few axons (Erlandson & Woodruff, 1982; Zheng, et al., 2008), rather than up to 20 small diameter axons in wildtype nerve Remak bundles (Harty & Monk, 2017). Notably, while neurofibromas rarely form, elevation of EGFR in wildtype SCs is sufficient to mimic this nerve-disruption phenotype (Ling, et al., 2005).

The *NF1* gene encodes neurofibromin, a GTPase activating protein that accelerates the hydrolysis of RAS-GTP to its inactive GDP-bound form downstream of EGFR (Simanshu, Nissley, & McCormick, 2017). In SCs, loss of neurofibromin causes increases in GTP-bound RAS (Kim et al., 1995; Sherman et al., 2000), and RAS-GTP stimulates the mitogen-activated protein kinase (MAPK) pathway and other downstream pathways. Loss of *NF1* also causes reduced levels of cyclic AMP (cAMP) in *Nf1* mutant mouse, *fly* and zebrafish (Hegedus, et al., 2007; Tong, et al., 2002; Wolman et al., 2014; Anastasaki and Gutmann, 2014). Whether cAMP deregulation occurs downstream of increased RAS-GTP is unclear. Neurofibromin shares homology with the yeast proteins Ira1 and Ira2, which are inhibitory regulators of the RAS–cAMP adenylyl cyclase pathway, but no evidence shows a direct role for NF1 or IRA proteins in direct regulation of cAMP in mammals (Ballester, et al., 1990; Martin, et al., 1990; Xu, et al., 1990). It is unclear how, or if, regulation of cAMP is relevant to neurofibroma initiation or growth but reducing cAMP drove formation of brain tumors in cells lacking *Nf1* (Warrington et al., 2010).

We sought to identify molecules that might affect neurofibroma development and regulate EGFR signaling expression in the SC lineage. We identified P2RY14 as a G-protein coupled receptor (GPCR) overexpressed in neurofibroma SCP and SC. P2RY14 is activated by extracellular UDP and UDP-sugars and signal through Gi to inhibit adenylate cyclase (AC), decreasing cAMP (Abbracchio & Ceruti, 2006; Conroy, Kindon, Kellam & Stocks, 2016). Intriguingly, P2RY14 regulates homeostasis of hematopoietic stem/progenitor cells (Cho et al., 2014). Also, satellite glial cells (SGCs) and SCs have been reported to express P2RY14 *in vitro* (Patritti-Cram, et al., 2021). Here we show that, *in vitro*, P2RY14 inhibition decreases mouse SCP self-renewal by modulation of cAMP and EGFR signaling. *In vivo*, P2RY14 knockout increased mouse survival and decreased tumor initiation and increasing cAMP in neurofibroma bearing decreased SC proliferation in neurofibromas. We suggest that targeting the P2RY14 receptor pathway could be relevant for treatment of NF1.

## Results

### P2RY14 is expressed in human neurofibroma Schwann cell precursors (SCP) and promotes SCP self-renewal *in vitro*

To characterize SCPs we used flow cytometry. We dissociated cells from human plexiform neurofibromas resected for therapeutic purposes from 3 neurofibroma patients. Cells were sorted into p75^+^/EGFR^-^ SCs and p75^+^/EGFR^+^ SCP-like tumor initiating cells. We performed gene expression analysis on these cells and found that *P2RY14* mRNA is elevated in p75^+^/EGFR^+^ SCP-like tumor initiating cells (**Figure 1A**). Western blot also showed expression in human SCs and neurofibroma SCPs, with 1.9-fold increase of P2RY14 protein in SCP-like cells (**Figure 1B**). To test if P2RY14^+^ SCP-like cells derived from human neurofibromas have altered ability to self-renew, we performed fluorescence activated cells sorting (FACS) and sorted SCP-like cells into p75^+^/EGFR^+^/P2RY14^-^ and p75^+^/EGFR^+^/P2RY14^+^ cells and plated them at low density to generate unattached spheres *in vitro* (**Figure 1C, D, E**). Unsorted neurofibroma cells rarely form SCP-like spheres. FACS analysis (of cells from 3 additional neurofibroma tumors) showed that on average p75^+^/EGFR^+^/P2RY14^-^ cells formed spheres at a frequency of 23.4%, while 64.8% of p75^+^/EGFR^+^/P2RY14^+^ cells formed spheres. The p75^+^/EGFR^+^/P2RY14^+^ cells maintained their significantly enhanced ability to form spheres *in vitro* for three passages (**Figures 1F, 1G**). Thus, P2RY14 is overexpressed in human neurofibroma SCs and SCPs *in vitro*, and marks SCP with the potential to self-renew *in vitro*.

**Figure 1:**
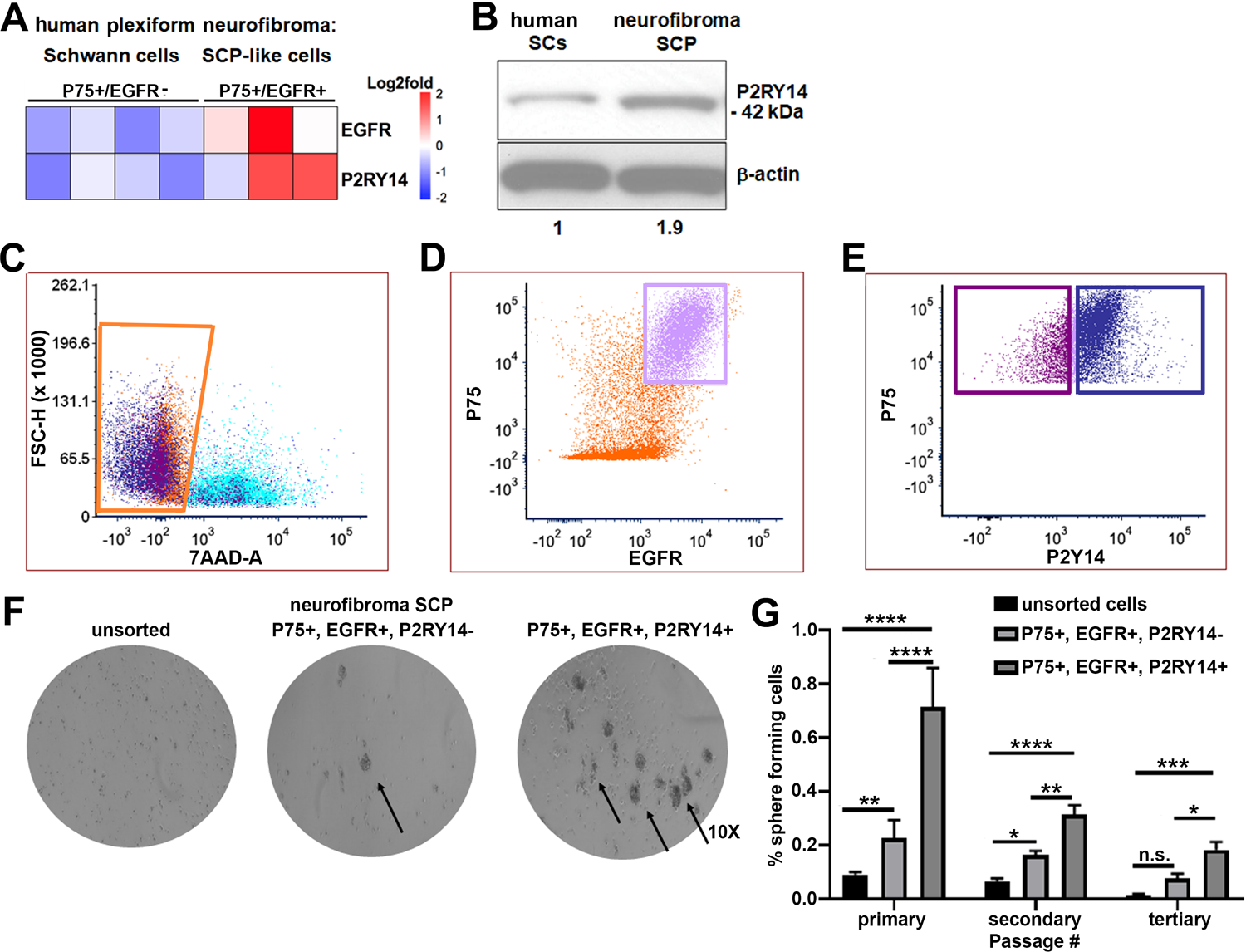
P2RY14 is expressed in human neurofibroma Schwann cell precursors and promotes SCP self-renewal *in vitro*. A) Microarray heatmap shows P2RY14 receptor expression in p75^+^/EGFR^+^ SCP-like tumor initiating cells derived from human plexiform neurofibroma tumor cells compared to p75^+^/EGFR^-^ SCP-like cells. B) Western blot of human Schwann cells and neurofibroma SCP shows the latter has a 1.9-fold increase in P2RY14 protein expression. C) Representative FACS plot shows live sorted human plexiform neurofibroma tumor cells. D) Representative FACS plot shows human plexiform neurofibroma tumor cells sorted into p75^+^/EGFR^+^ SCP-like tumor initiating cells (pink square). E) Representative FACS plot shows p75^+^/EGFR+ SCP-like tumor initiating cells further sorted into p75^+^/EGFR^+^/P2RY14^-^ (left, purple square) and P75^+^/EGFR^+^/P2RY14^+^ (right, blue square). F) Photomicrographs of human neurofibromas dissociated using FACS to yield: p75^+^/EGFR^+^/P2RY14^-^ and P75^+^/EGFR^+^/P2RY14^+^ cells. G) Quantification of unsorted, p75^+^/EGFR^+^/P2RY14^-^ and P75^+^/EGFR^+^/P2RY14^+^ cells plated in sphere medium. (n=3; two-way ANOVA; primary: **p=0.0057, ****p<0.0001; secondary: *p=0.0487,**p<0.0024, ****p<0.0001; tertiary:*p=0.0321, ***p=0.0006).

### Mouse *Nf1* mutant SCPs use P2RY14 signaling through Gi to regulate self-renewal

We verified P2RY14 expression in cultured SCP spheres from wild-type (*WT*) and *Nf1-/-* mouse embryos. P2RY14 protein was slightly elevated in *Nf1^-/-^* SCPs (**Figure 2A**). P2RY14 signaling is relevant in this setting, as pharmacological inhibition of P2RY14 with the highly selective P2RY14 inhibitor PPTN (4-[4-(4-Piperidinyl)phenyl]-7-[4-(trifluoromethyl)phenyl]-2-naphthalenecarboxylic acid hydrochloride) (Robichaud et al., 2011) reduced the percentage of *Nf1^-/-^* mouse SCPs that formed spheres, but had little effect on wild-type SCPs, consistent with a role in *Nf1* mutant SCP self-renewal (**Figure 2B**). Photographs of spheres are shown in **Supplemental figure 1A**; this experiment was repeated in 3 biological replicates with similar results. Dose response analysis confirmed the optimal concentration of the P2RY14 inhibitor (PPTN) is 300nM; as 500nM PPTN was toxic (**Supplemental figures 2A-2F**).

**Figure 2:**
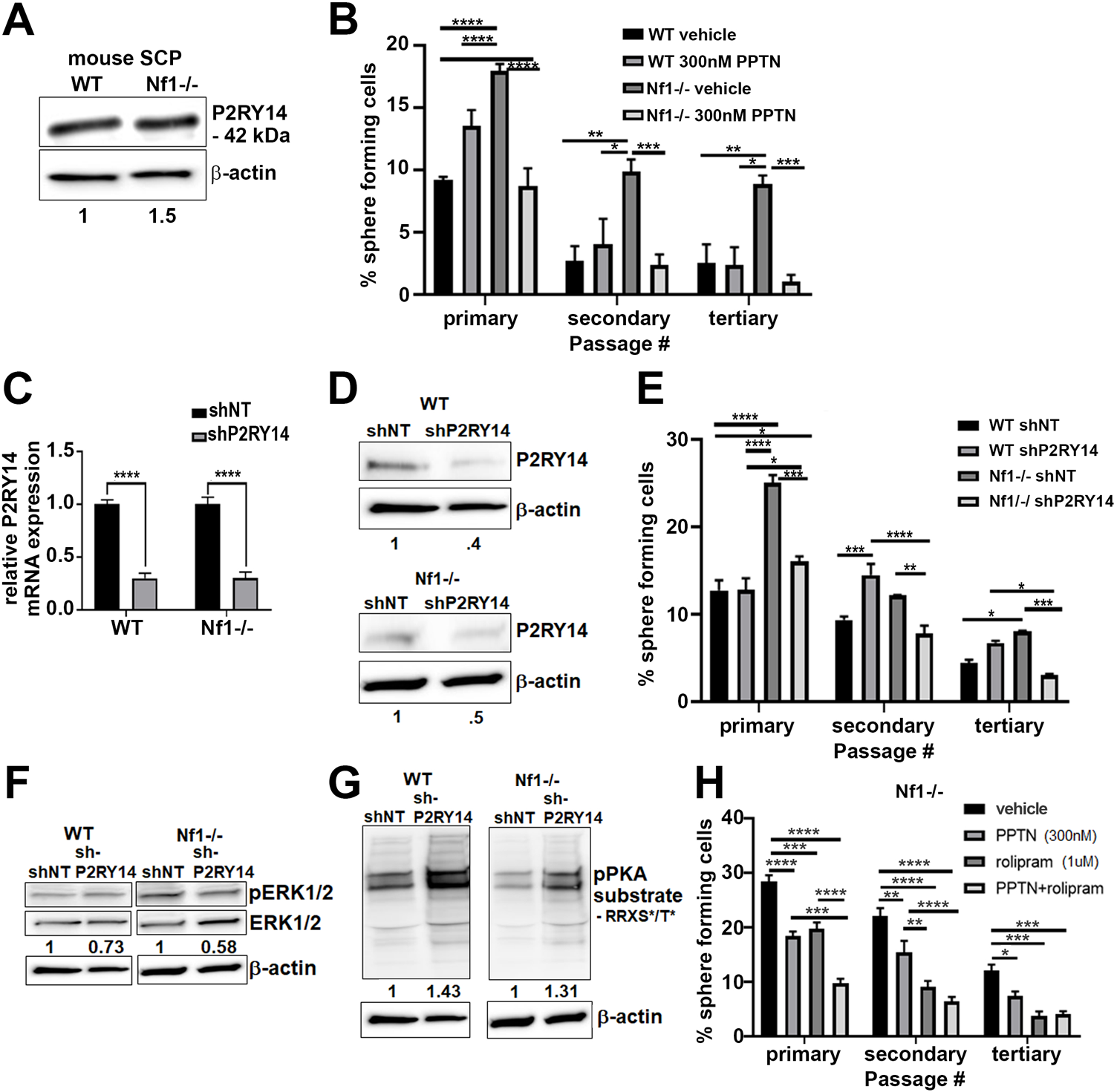
Mouse *Nf1* mutant SCPs use P2RY14 signaling through Gi to regulate self-renewal. A) Western blot shows Nf1^-/-^ SCPs have 1.5-fold change increase in P2RY14 expression. B) Quantification of percent of sphere forming cells in mouse *WT* and *Nf1^-/-^* SCPs treated with the selective P2RY14 inhibitor (300nM PPTN) (primary, secondary, tertiary passage) (n=3; two-way ANOVA; primary: *p=0.0375, ***p=0.0001, ****p<0.0001; secondary: *p=0.0101, **p=0.0015, ***p=0.0009; tertiary: *p=0.0101, **p=0.0050, ***p=0.0005). C) P2RY14 mRNA expression in *WT* and *Nf1^-/-^* E12.5 mouse SCP treated with shnon-target (shNT) control and shP2RY14 (****p<0.0001). D) Western blot of *WT* and *Nf1^-/-^* SCPs treated with shNT and shP2RY14 showing P2RY14 knockdown. *WT* shP2RY14 show a 0.4-fold decrease of P2RY14 protein compared to *WT* shNT; and Nf1^-/-^ shP2RY14 show a 0.5-fold decrease compared to *Nf1^-/-^*. E) Quantification of percent of sphere forming cells in mouse *WT* and *Nf1^-/-^* SCPs treated with shNT and shP2RY14 (n=3; two-way ANOVA; primary: *p=0.0288, ****p<0.0001; secondary: **p=0.0029,***p=0.0005, ****p<0.0001; tertiary: *p=0.0154, ***p=0.0007). F) Western blot of *WT* and *Nf1^-/-^* SC spheres shows 0.73-fold decrease of pERK in *WT* cells and 0.58-fold pERK decrease in *Nf1^-/-^* cells after P2RY14 knockdown. G) Western blot of *WT* and *Nf1^-/-^* SC spheres shows changes in pPKA substrate phosphorylation after shP2RY14 knockdown. *WT* shP2RY14 show a 1.43-fold increase in pPKA after P2RY14 knockdown; *Nf1-/-* cells have a 1.31-fold increase in pPKA expression after P2RY14 knockdown. H) Quantification of percent of sphere forming cells in *Nf1^-/-^* mouse SCPs treated with 1uM rolipram or 300nM PPTN (n=3; two-way ANOVA; primary: ***p=0.0002, ****p<0.0001; secondary **p=0.0030, ****p<0.0001; tertiary: *p=0.0476, ***p=0.0004).

To confirm these results, we silenced P2RY14 gene expression using short-hairpin RNAs (shRNAs) targeting P2RY14. *WT* and *Nf1^-/-^* mouse SCPs were treated with non-target control (shNT) or shRNA P2RY14 (shP2RY14). shP2RY14 treated cells showed reduced *P2RY14* mRNA (**Figure 2C**) and P2RY14 protein (**Figure 2D**). Sphere formation was significantly decreased in *Nf1^-/-^* SCPs but not wild-type treated with *P2RY14* shRNA, and this phenotype also persisted for 3 passages (**Figure 2E**). Photomicrographs are shown for one shRNA (sh09) (**Supplemental figure 1B**); but the experiment was repeated with 2 additional P2RY14 shRNAs in 3 biological replicates each, with similar results (**Supplemental figure 3A-3D**). To delineate signaling pathways affected by P2RY14 shRNA in SCPs, we analyzed cell lysates in western blots. We found that shP2RY14 caused a slight decrease in pERK, a read-out of RAS/MAPK signaling, which we know to be elevated in *Nf1-/-* cells (**Figure 2F**). We analyzed cAMP-dependent protein kinase (PKA) substrate phosphorylation (using anti p-PKA substrate antibody) as an indirect read out of cAMP levels in cells. p-PKA substrate phosphorylation increased in *WT* and *Nf1-/-* SCPs shP2RY14 treated cells. Interestingly, knockdown of P2RY14 in *Nf1^-/-^* SCPs increased levels of pPKA, similar to those in WT shNT levels (**Figure 2G**).

**Figure 3:**
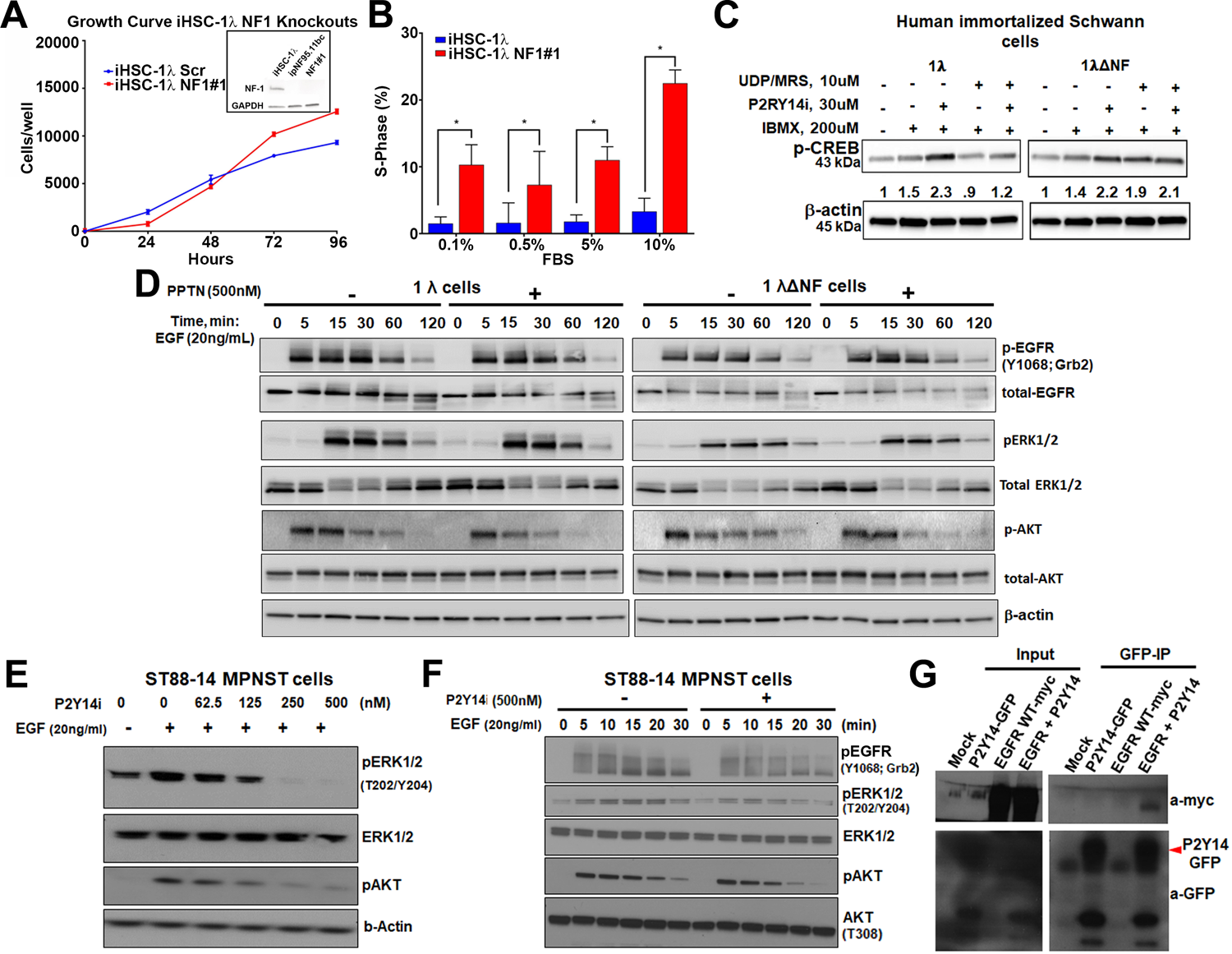
P2RY14 decreases EGFR phosphorylation suppressing EGFR-stimulated pERK and pAKT. A) Growth curve of iHSC1λ and iHSC1λ NF1#1 (1λ**Δ**NF). Top right inset shows western blot confirming NF1 protein expression in iHSC 1λ (positive control), ipNF95.11bc SCs (negative control) and iHSC 1λ NF1#1 (1λ**Δ**NF) (experimental). B) EdU incorporation assay in iHSC1λ and iHSC1λ NF1#1 (1λ**Δ**NF) (*p<0.001). C) 1λ and 1λ ΔNF1 treated with a potent 10uM UDP analog (P2RY14 specific agonist MRS2690), 500nM of the P2RY14 inhibitor (PPTN) and 200uM of IBMX (n=3). Western blot show changes in p-CREB expression upon treatment. D) 1λ and 1λΔNF1 cells treated with the P2RY14 inhibitor (PPTN; 500nM) at different timepoints (0,5,15,30,60,120 minutes) after EGF stimulation and western blot shows changes in pEGFR-Grb2 (Y1068), total EGFR, pERK1/2, total ERK1/2, p-AKT, total AKT and β-actin. E) Western blot shows pERK1/2, ERK1/2 and pAKT changes in ST88-14 cells treated with different concentrations of the P2RY14 inhibitor (PPTN) (0, 62.5nM, 125nM, 250nM and 500nM of PPTN) and 20ng/mL of EGF. F) Western blot shows ST88-14 cells treated with the P2RY14 inhibitor (PPTN; 500nM) and EGF (20ng/mL) and shows changes in EGFR phosphorylation at pEGFR-Grb2 (Y1068) binding site and changes in pERK and pAKT. G) ST88-14 cells transfected with P2RY14-GFP and/or EGFR-myc tagged proteins and immunoprecipitated EGFR using an anti-myc antibody shows the formation of a P2RY14 complex with EGFR.

To understand if P2RY14-mediated changes in cAMP signaling affect SCP self-renewal, we treated *WT* and *Nf1^-/-^* SCPs with the specific phosphodiesterase-4 (PDE4) inhibitor, rolipram. Rolipram blocks degradation of cAMP by PDE4, increasing intracellular levels of cAMP (MacKenzie & Houslay, 2000). Treatment with 1uM rolipram or 300nM of the P2RY14 inhibitor (PPTN) decreased *Nf1^-/-^* SCP self-renewal; the combination showed additional effect, largely at early passage (**Figure 2H**). Photographs of these experiments are shown in **Supplemental Figure 1C**. These results support the idea that the elevated self-renewal in *Nf1^-/-^* SCP is due, at least in part, to P2RY14 Gi-mediated changes in cAMP.

### P2RY14 inhibition increases cAMP and decreases EGFR activation and downstream signaling

To test if the relationship between P2RY14, cAMP and NF1 found in SCPs also exists in SCs, we turned to cell lines that enable biochemical analysis and used human immortalized SCs (iHSCs) (Li et al., 2016). We used lentivirus infection and the CRISPR/Cas9 system to generate 1λΔNF1 (NF1 deficient iHSCs). Pooled puromycin resistant cells were evaluated for NF1 growth over time and NF1 expression (**Figure 3A**). Increased proliferation was confirmed by increased EdU incorporation of 1λ ΔNF1 cells (**Figure 3B**). To confirm effects of P2RY14 in this setting, we treated 1λ iHSCs (wildtype for NF1) and 1λΔNF1 (NF1 deficient cells) with 10uM of a potent P2RY14-specific UDP agonist (MRS2690), 500nM of the P2RY14 inhibitor (PPTN), and 200uM IBMX (an inhibitor of PDEs). We measured phosphorylated cAMP response element binding protein (pCREB) as an indirect readout of cAMP. In 1λ iHSCs, addition of the P2RY14 inhibitor increased pCREB expression, as expected, since inhibition of P2RY14 releases Gi mediated inhibition of adenylate cyclase (AC), increasing intracellular levels of cAMP. In the absence of the P2RY14 inhibitor but in the presence of the potent P2RY14-specific UDP agonist (MRS2690), we observed a decreased in pCREB expression, confirming that upon activation by UDP, P2RY14 acts through Gi to inhibit AC and decrease cAMP (**Figure 3C**). In 1λΔNF1 cells, addition of the P2RY14 inhibitor similarly increased pCREB expression. However, while in wild type cells activation of P2RY14 with MRS2690 in 1λ cells decreased pCREB expression, in 1λΔNF1 cells, the increased expression was maintained upon addition of the potent P2RY14-specific UDP agonist (**Figure 3C**), suggesting that activation of P2RY14 resulted in sustained pCREB, even in the presence of agonist (**Figure 3C**). To test if NF1 regulates cAMP directly we tested non-immortalized mouse primary SCs. *WT* mouse primary SCs showed elevated levels of basal cAMP compared to isogenic *Nf1-/-* SCs. Activation of AC via forskolin stimulation increased cAMP in *WT* SCs, but to a lesser extent in *Nf1*-/-SCs (**Supplemental figure 3E**). Based on these results, in both immortalized SCs and mouse primary SCs, NF1 alters cAMP signaling.

Given that EGFR and P2RY14 similarly modulate SCP self-renewal, we hypothesized that P2RY14 inhibition might affect EGFR downstream signaling. To test this idea, we treated 1λ and 1λΔNF1 with epidermal growth factor (EGF), with and without the P2RY14 inhibitor (PPTN). We monitored EGFR-phosphorylation at the Grb2 activation site to measure EGFR activation, and examined downstream signaling with pERK1/2 and phosphorylation of Akt at the threonine 308 (T308) activation site (Vincent, et al., 2011). In 1λΔNF1 cells, P2RY14 inhibition reduced phosphorylation of EGFR at the Grb2 activation site and shortened the duration of p-AKT signaling (**Figure 3D**; Supplemental figure 4). Similar results were seen in the *NF1* deficient MPNST cell line, ST88-14 (**Figure 3E-3F**). To explain these results, we hypothesized that P2RY14 might form a complex with EGFR. To test this idea, we transfected ST88-14 cells with P2RY14-GFP and/or EGFR-myc tagged proteins. Then, we immunoprecipitated EGFR using an anti-myc antibody and detected P2RY14 in a complex with anti-GFP EGFR (**Figure 3G**). **P2RY14 is expressed *in vivo* in mouse SCPs and SCs.** In the *Nf1fl/fl;Dhh+* neurofibroma mouse model, Cre recombinase is expressed from Desert Hedgehog (Dhh) regulatory sequences to effect recombination of the *Nf1 fl/fl* allele, resulting in loss of both *Nf1* alleles in developing SCPs at embryonic day 12.5 (Wu et al., 2008). *Nf1fl/fl;Dhh+* mice develop paraspinal neurofibromas that have loss of axon-Schwann cell interaction in Remak bundles, mast cell and macrophage accumulation, and nerve fibrosis, all characteristics of human plexiform neurofibromas (Wu et al., 2008; Liao, et al., 2018; Fletcher, et al., 2019(a); Fletcher, et al., 2019 (b)). In this setting, plexiform neurofibromas are present by 4-months of age, and neurofibromas enlarge as the mice age, ultimately compressing the spinal cord causing paralysis and thus, necessitating sacrifice (Wu et al., 2008). We bred *P2RY14^-/-^* mice (Meister et al., 2014) to generate *P2RY14****^-/-^****;Nf1fl/fl;Dhh+* and *P2RY14****^+/-^****^;^Nf1fl/fl;Dhh+* littermates, and *Nf1 fl/fl Dhh+* controls (**Figure 4A**). Genotyping and western blotting verified reduced P2RY14 in *P2RY14^-/-^;Nf1fl/fl;Dhh+* sciatic nerve and neurofibroma tumors compared to *Nf1fl/fl Dhh+* controls (**Figure 4B & 4C**). In the *P2RY14^-/-^* mice, most of the coding region of P2RY14 is replaced by a β-galactosidase cassette, so that where the P2RY14 gene is expressed, β-galactosidase is detectable (Meister et al., 2014). We visualized P2RY14 in SOX-10 positive SCPs in the spinal cord dorsal and ventral roots (VR) and in the dorsal root ganglion (DRG) of embryonic 12.5 mice (**Figure 4D & 4E**). *P2RY14^-/-^;Nf1fl/fl;Dhh+* SCs in 7-month old mice were also positive for β-galactosidase staining (**Figure 4F**). CNPase (2’, 3’ cyclic nucleotide 3’ phosphodiesterase), a known SC marker, co-localized with β-galactosidase staining in the sciatic nerve (**Figure 4F, inset**). P2RY14 antibody staining confirmed P2RY14 expression in myelinating SC in mouse *WT* nerve and in *Nf1fl/fl;Dhh+* neurofibromas (**Figure 4G**).

**Figure 4:**
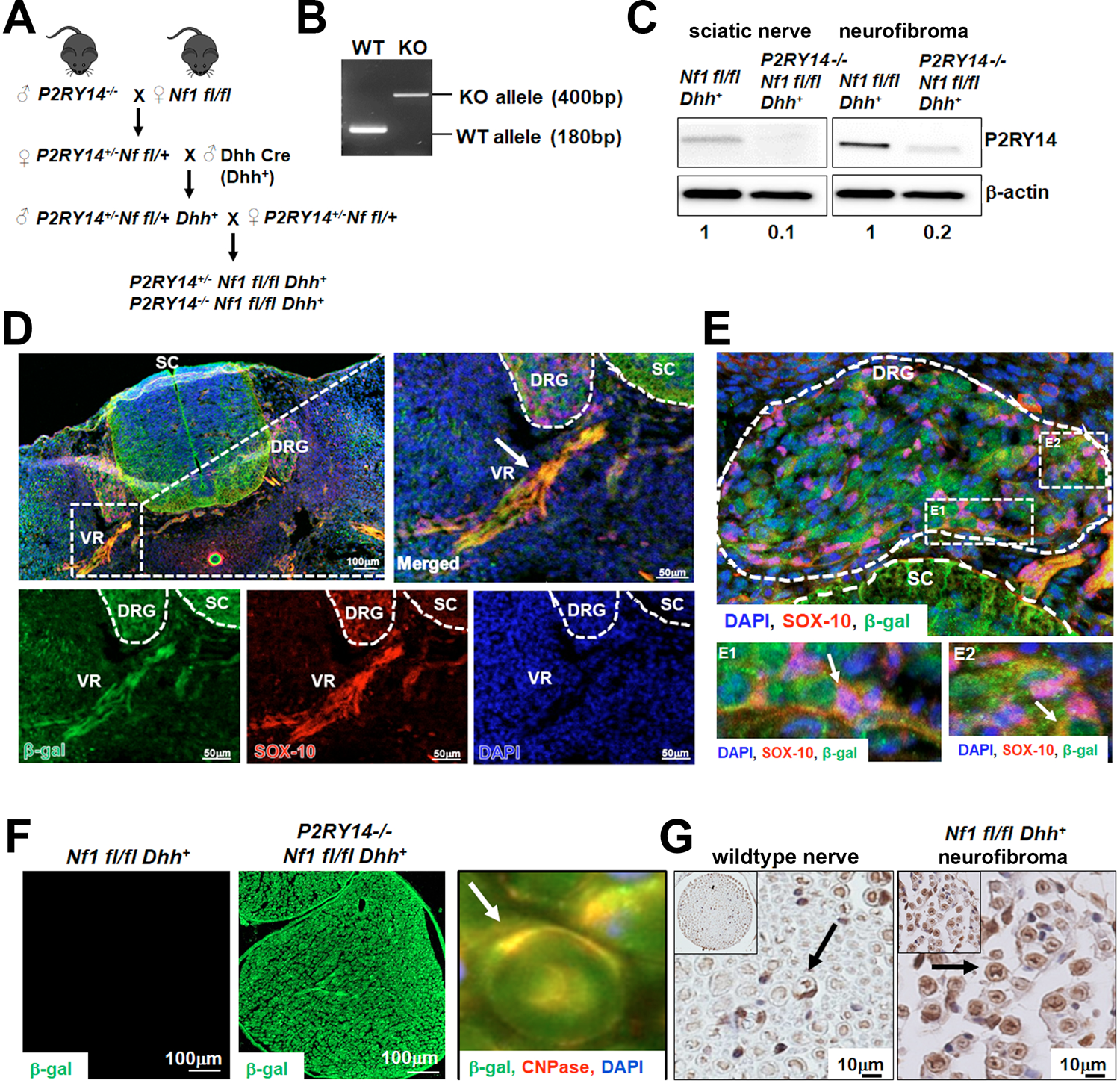
P2RY14 loss in mouse model of NF1 rescues reduced PKA mediated phosphorylation. A) Schematic of generation of neurofibroma mice breeding of *P2RY14^-/-^* mice with *Nf fl/fl* mice to obtain *P2RY14^+/-^;Nf1fl/fl;Dhh+* and *P2RY14^-/-^;Nf1fl/fl*;*Dhh+* after several crosses. B) Genotyping confirmation of wildtype (WT) and P2RY14 knockout (KO) alleles. C) Western blot of sciatic nerve and neurofibroma tumors of *Nf1fl/fl;Dhh+* and *P2RY14^-/-^;Nf1fl/fl*;*Dhh+* mice show decrease in P2RY14 expression upon P2RY14 knockdown (1 to 0.1-fold decrease in sciatic nerve and 1 to 0.2-fold decrease in neurofibroma tissue). D) Spinal cord (SC) immunofluorescent staining of mouse embryos at E12.5 shows P2RY14 expression (β-galactosidase) co-localization with SOX-10 SCs at dorsal and ventral roots (VR). E) Spinal cord (SC) immunofluorescent staining of mouse embryos at E12.5 shows P2RY14 expression (β-galactosidase) co-localization with SOX-10 SCs at dorsal root ganglion (DRG). E1 and E2 insets show zoomed picture of DRG. F) Immunofluorescent staining of 7-month-old mouse sciatic nerve shows β-galactosidase positive staining as a confirmation of P2RY14 knock-in; co-labeling of β-galactosidase and CNPase shows that P2RY14 co-localizes with SCs (inset). G) DAB staining of 7-month old *WT* nerve and *Nf1fl/fl;Dhh+* mouse neurofibromas (DAB staining: brown (P2RY14 positive cells) blue (cell nuclei)).

### P2RY14 deletion in mouse model of neurofibroma increases survival, delays neurofibroma initiation, and rescues SC Remak bundle disruption

Kaplan Meier survival analysis showed that *P2ry14^-/-^;Nf1fl/fl;Dhh+* have a significant survival advantage compared to neurofibroma-bearing mice (*Nf1fl/fl;Dhh+*; p=0.0256). In fact, *P2RY14^-/-^;Nf1fl/fl;Dhh+* did not differ significantly from non-neurofibroma bearing *Nf1fl/+;Dhh^+^* control littermates (p=0.1367) (**Figure 5A**). To test if *P2RY*14 deletion plays a role in neurofibroma initiation, we analyzed *Nf1fl/fl;Dhh+* and *P2RY14 ^-/-^;Nf1fl/fl;Dhh+* mice to 4-months of age, when tumor are first detectable. Gross dissection of spinal cords from these mice showed that *P2RY14^-/-^;Nf1fl/fl;Dhh+* have decreased neurofibroma number, consistent with a role in tumor initiation, and only slightly reduced neurofibroma diameter (**Supplemental figure 5A, 5B, 5C**). Ki67 and H&E staining of neurofibromas in these 4-month-old mice showed no evident changes between genotypes in cell proliferation or cell morphology (**Supplemental figure 5D-5F**). To determine if the effects of P2RY14 persist, we aged *Nf1fl/fl;Dhh+* and *P2RY14^-/-^;Nf1fl/fl;Dhh+* mice to 7-months. *P2RY14^-/-^;Nf1fl/fl;Dhh+* mice also showed significantly fewer neurofibromas compressing the spinal cord on gross dissection (**Figure 5B, 5C**), and tumor diameter was reduced to a lesser degree (**Figure 5D**). By this age, *P2RY*14 loss decreased SC proliferation in neurofibroma tissue sections, at least in part in CNPase^+^ SCs (**Figure 5E-5F**; **Supplemental Figure 5G**) yet H&E staining retained characteristic cell morphology (**Supplemental figure 5H**). Confirming potential relevance of P2RY14 in SCs, P2RY14 protein was detected in membranes of myelinating SCs in human neurofibromas tissue sections (**Figure 5G**).

**Figure 5:**
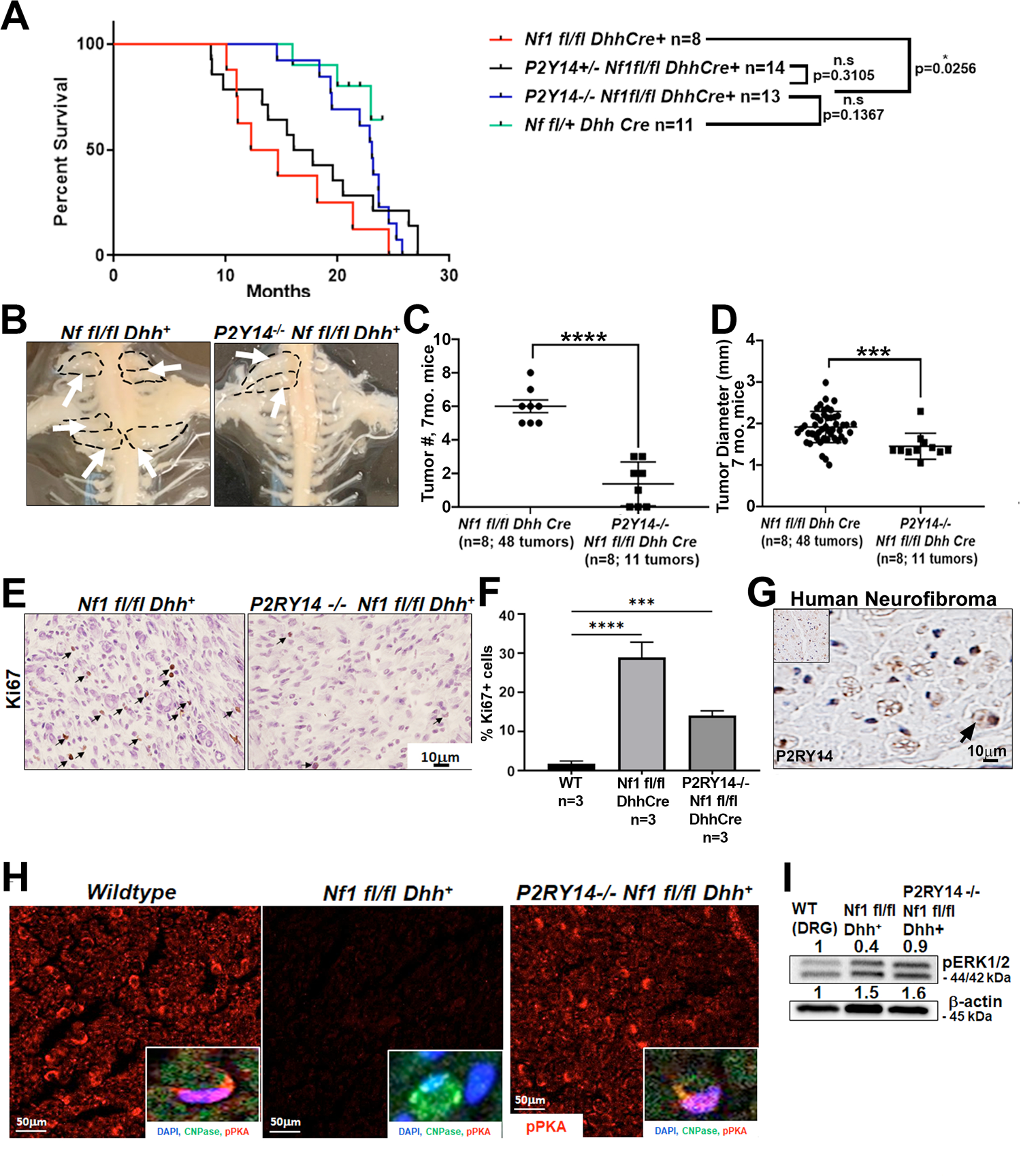
P2RY14 deletion in mouse model of neurofibroma increases survival and delays neurofibroma initiation. A) Kaplan Meier survival plot of *Nf1fl/fl;Dhh+* (red line; n=8); *P2RY14^+/-^;Nf1fl/fl;Dhh+* (black line, n=14); *P2RY14^-/-^;Nf1fl/fl*;*Dhh+* (blue line; n=13); *Nf1fl/+* control (green line, n=11) (*p=0.0256). B) Representative image of gross dissection of *Nf1fl/fl*;*Dhh+* and *P2RY14^-/-^;Nf1fl/fl*;*Dhh+* mice at 7-months of age. C) Neurofibroma tumor number quantification at 7-months of age (unpaired *t* test ****p<0.0001). D) Neurofibroma diameter quantification at 7-months (unpaired *t* test ***p=0.0004) (for figures C & D: *Nf1fl/fl*;*Dhh^+^* n=8 mice, 48 neurofibroma tumors; *P2RY14-/-;Nf1fl/fl*;*Dhh+* n=8 mice, 11 neurofibroma tumors). E) Ki67 staining of mouse DRG and neurofibroma tissue at 7-months of age. F) Quantification of Ki67+ cells in mouse DRG and neurofibroma tissue at 7-months of age (One-way ANOVA; multiple comparisons ***p=0.0008; ****p<0.0001). G) Immunohistochemistry of human neurofibroma shows P2RY14 expression (DAB staining: brown (P2RY14 positive cells) blue (cell nuclei)). H) anti p-PKA substrate staining in *WT*, *Nf1fl/fl;Dhh+* and *P2RY14^-/-^;Nf1fl/fl,Dhh+* mice. p-PKA substrate phosphorylation labeling co-localized with CNPase SC marker (insets). I) Western blot of tissue lysates of *WT* (DRG), *Nf1fl/fl;Dhh+* (neurofibroma tumors) and *P2RY14^-/-^;Nf1fl/fl*;*Dhh+* (neurofibroma tumors) show *Nf1fl/fl;Dhh+* neurofibromas have 1.5-fold increased p-ERK expression compared to *WT* DRGs but, p-ERK expression between *Nf1fl/fl;Dhh+* (1.5-fold) and *P2RY14*-/-*;Nf1fl/fl*;*Dhh+* (1.6-fold) remained relatively unchanged.

To determine if cAMP is affected by loss of *P2RY14* in tumors, we stained with anti p-PKA substrate antibody. *Nf1fl/fl;Dhh+* nerves showed a decrease in p-PKA substrate phosphorylation versus wild-type mice. Remarkably, this decrease was reversed in *P2RY14^-/-^;Nf1fl/fl;Dhh+* nerves (**Figure 5H**). pPKA labeling co-localized with CNPase, suggesting that the changes in p-PKA substrate phosphorylation expression, are at least in part, in nerve SCs (**Figure 5H, insets**). Conversely, p-ERK was modestly increased by loss of *P2RY*14 in *Nf1fl/fl;Dhh+* mice (**Figure 5I**). Based on these results, we conclude that *P2RY*14 deletion *in vivo* in neurofibroma mice increases mouse survival and delays neurofibroma initiation, with lesser effects on SC proliferation. Importantly, deletion of P2RY14 also rescues reduced PKA mediated phosphorylation in *Nf1fl/fl;Dhh+* SCs.

To study an additional SC phenotype in these mice, we examined the effects of *P2RY*14 loss on nerve disruption phenotype using electron microscopy. At 4-months-old *Nf1fl/fl;Dhh+* mice already had disrupted Remak bundles (reduced numbers of axons ensheathed by individual SCs); this phenotype was rescued in *P2RY14^-/-^;Nf1fl/fl;Dhh+* mice (**Supplemental figure 6A, 6C**). By 7-months, *Nf1fl/fl;Dhh+* nerves were even more severely disrupted, and rescued to near wild-type levels by *P2RY14* loss (**Supplemental figure 6B, 6D**). We conclude that genetic knockout of P2RY14 in neurofibroma bearing mice rescues the nerve Remak bundles defects characteristic of neurofibroma bearing mice.

**Figure 6:**
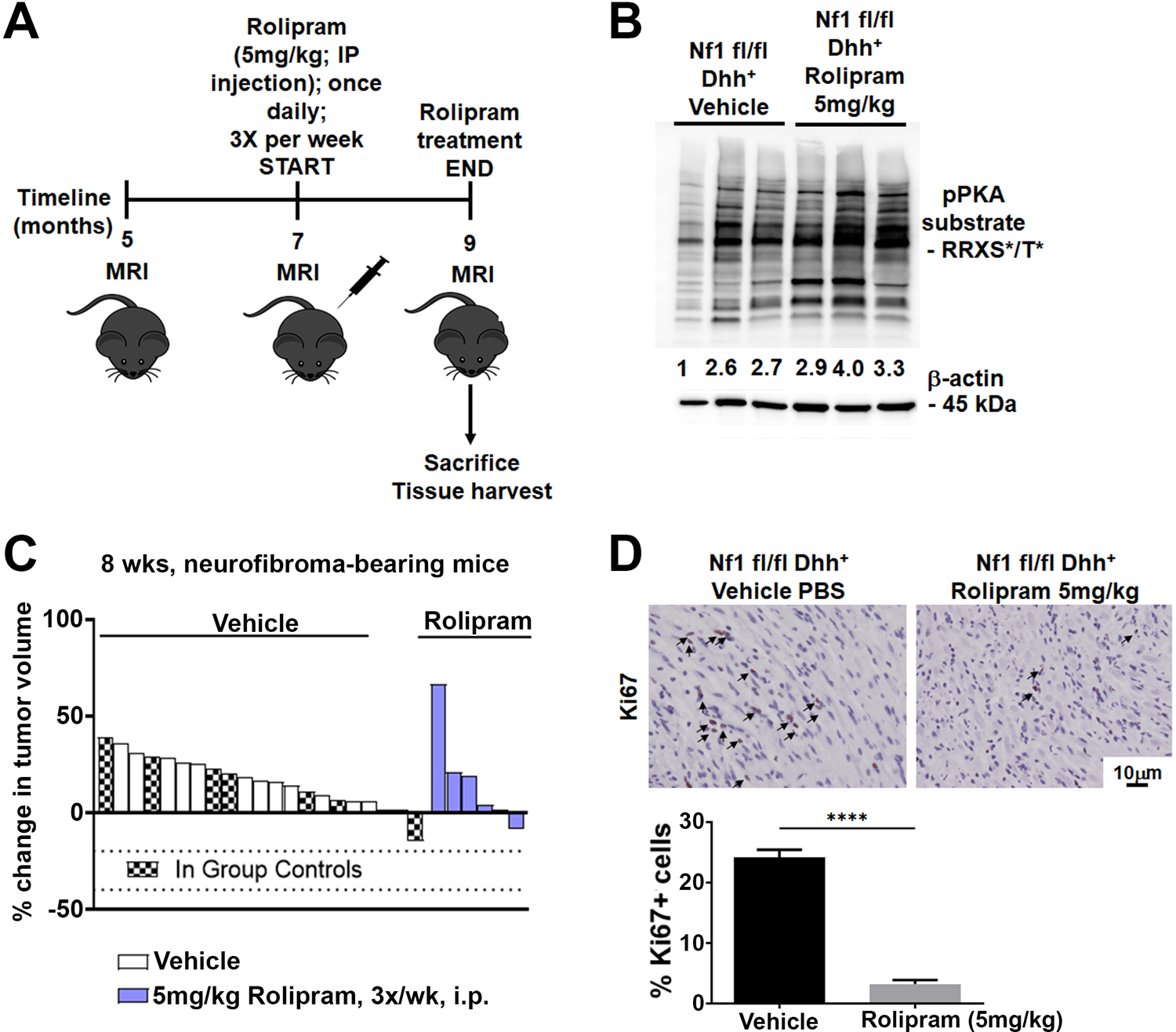
cAMP increase in neurofibroma bearing mice via rolipram treatment decreases SC proliferation in neurofibromas. A) Rolipram drug treatment experimental design. B) Tumor lysates of vehicle and rolipram treated *Nf1fl/fl*;*Dhh+* mice show changes in p-PKA substrate. C) Volumetric analysis of neurofibroma volumes in *Nf1fl/fl*;*Dhh+* mice treated with 5mg/kg rolipram (Unpaired t-test: p=0.9397). D) Ki67 staining at 9-months of age in vehicle and rolipram treated mice. Inset shows quantification of Ki67+ cells in vehicle treated versus rolipram treated mice (unpaired t-test: ****p<0.0001; n=3).

### Increasing cAMP in neurofibroma bearing mice by means of rolipram treatment decreases SC proliferation in neurofibromas

To test if cell proliferation in *Nf1fl/fl;Dhh+* neurofibromas is affected by changes in cAMP levels, *Nf1fl/fl;Dhh+* mice were treated with vehicle or 5mg/kg rolipram (**Figure 6A**). All mice survived rolipram treatment without significant weight loss. Rolipram treated neurofibroma mice tumors lysates showed increases in p-PKA substrate phosphorylation (**Figure 6B**). Neurofibroma volume in *Nf1fl/fl;Dhh+* mice treated with rolipram did not significantly differ from controls (**Figure 6C**), consistent with the idea that P2RY14 plays a major role in tumor initiation and a minor role in neurofibroma growth. However, cell proliferation (Ki67 staining) in tissue sections showed that rolipram treated mice have decreased cell proliferation *in vivo* (**Figure 6D**), although they did not show a significant decrease in tumor volume. These results are consistent with the idea that reduction in neurofibroma cell proliferation is at least in part mediated by cAMP, as increases in cAMP by addition of rolipram, like loss of P2RY14, decreases neurofibroma cell proliferation. Together, these studies show that the G-protein P2RY14 is a key regulator of neurofibroma initiation, and that P2RY14 modulates EGFR-driven self-renewal, and SCP/SC cAMP signaling.

## Discussion

GPCR-mediated regulation of cAMP occurs upon *NF1* loss in mammals (Dasgupta, Dugan, & Gutmann, 2003; Deraredj Nadim, et al., 2016), fish, and dropsophila (Tong, et al., 2002; Wolman et al., 2014). In these systems, PACAP receptors and serotonin receptors have been identified as GPCRs that act upstream of NF1 (Anastasaki & Gutmann, 2014; Deraredj Nadim et al., 2016). However, the potential role of cAMP in neurofibroma remains unclear. Targeting GPCR signaling has been suggested as a potential therapeutic option to treat NF1, so exploring the relevance of this pathway to peripheral nerve tumors is important.

We found that human neurofibroma derived-SCP-like cells sorted for G-protein coupled receptor P2RY14 expression have increased self-renewal potential. Adult peripheral nerves do not contain a stem cell population (Stierli, et al., 2018). Therefore, neurofibroma SCP-like cells may result from persistence of immature cells and/or from de-differentiation of mutant Schwann cells. In either case, neurofibroma also contain cells that also express P2RY14. Our findings are entirely consistent with findings that hematopoietic stem/progenitor cells marked by P2RY14 stimulate self-renewal (Lee, et al., 2003; Cho, et al., 2014; Holmfeldt, et al., 2016).

Purinergic receptor P2RY14 is activated by UDP and by the nucleotide sugars UDP-glucose, UDP-galactose, UDP-glucuronic acid and UDP-N-acetylglucosamine (Chambers et al., 2000; Moore et al. 2003; Abbracchio et al., 2003; Carter et al., 2009; Conroy et al., 2016). *NF1* mutant cells may release one or more of these ligands, because SCP self-renewal in serum-free medium was reduced by pharmacologic P2RY14 inhibition, and by shRNA targeting P2RY14, in the absence of added UDP or UDP-sugars. The small increase in P2RY14 protein in mutant cells may contribute to increasing signaling downstream of the GPCR, and/or *Nf1* deficient cells may release more UDP-sugars than wild type cells. While it is difficult to measure UDP levels in the extracellular milieu without causing cell damage and concomitant release UDP sugars (Lazarowki & Harden, 2015), it will be of interest to measure UDP and UDP-sugars both in the neurofibroma extracellular milieu, and in SCP and SC culture medium.

UDP, UTP, and other nucleotide sugars including UDP-glucose are present at high levels in tumor cells, and released from cells in tumors and injury sites, where they act as danger signals that trigger inflammatory responses (Eigenbrodt, Reinacher, Scheefers-Borchel, Scheefers, & Friis, 1992; Skelton, Cooper, Murphy, & Platt, 2003). We focused on the role of P2RY14 in SCs and SCPs, because most of the P2RY14 expression in peripheral nerve neurofibromas is in CNPase^+^ myelinating Schwann cells, based on our use of a β-galactosidase reporter and antibody staining. However, P2RY14 is also expressed in immune cells and in small sensory neurons and larger diameter sensory neurons (Skelton, Cooper, Murphy, & Platt, 2003; Müller, et al., 2005; Scrivens and Dickenson., 2005; 2006; Malin & Molliver, 2010). In our studies, because we used a global knockout, we cannot exclude the idea that some of the observed *in vivo* effects are SCP-independent. The recent description of a conditional knockout of P2RY14 should enable additional studies (Battistone, et al., 2020).

In *NF1* mutant cells, pharmacological inhibition of P2RY14 also decreased activation of EGFR/ pAKT(T308) activation site in SCPs, as well as in iHSC and MPNST cells. Consistent with the idea that GPCR-receptor tyrosine kinase crosstalk occurs in the absence of *NF1*, P2RY14 loss rescued the Remak bundle disruption which is a key feature of neurofibroma SCs and is mimicked by transgenic expression of EGFR expression in Schwann cells. Cross-talk between GPCRs and EGFR (or the related receptor ErbB2) can occur through physical interaction. This may occur in *NF1*-deficient cells, as we found that P2RY14 and EGFR can form a complex after overexpression in MPNST cells. Whether such a complex facilitates EGFR signaling in SCP and SCs remains to be determined.

Whether the decreased levels of cAMP in *NF1* deficient cells requires upstream RAS signaling is unclear (reviewed in Bergoug et al., 2020). In *NF1* mutant neurons, RAS activation of atypical protein kinase C-zeta causes GRK2-driven Gαs/AC inactivation downstream of the PACAP receptor (Anastasaki & Gutmann, 2014). In fly, some investigators proposed direct regulation of AC by loss of NF1 (Tong, Hannan, Zhu, Bernards, & Zhong, 2002), but others propose that regulation is indirect (Walker & Bernards, 2014). In addition, PKA phosphorylates neurofibromin and inhibits its GAP activity (Feng, et al., 2004), suggesting potential negative feedback between NF1/RAS and PKA in wildtype cells. Furthermore, in some cell types, downstream of P2RY14/Gi/o, Ca2^+^ mobilization can result in release of factors such as IL-18, which could activate RAS indirectly (Moore et al., 2003; Arase et al., 2009). All these mechanisms are candidates to explore for relevance to P2RYR14 signaling in SCPs and SCs.

Genetic knockdown of P2RY14 increased levels of cAMP-dependent protein kinase (PKA)-mediated phosphorylation in SCP *in vitro* and SC *in vivo*, and *NF1^-/-^* SCPs treated with rolipram to increase cAMP showed decreased SCP self-renewal. *In vivo,* neurofibroma bearing mice treated with rolipram showed increases in pPKA expression. These results implicate cAMP modulation as a critical P2RY14 effector in both cell types. Consistent with these findings, in wildtype cells, P2RY14/Gi/o-mediated decreases in cAMP affect chemotaxis hematopoietic of stem/progenitor cells (Lee et al., 2003; Scrivens and Dickenson, 2005). Supporting a role for P2RY14-Gi-driven suppression of cAMP levels in SCP/SC, p-PKA substrate staining was reduced in *Nf1* mutant nerves and elevated by additional loss of *P2RY14*. Importantly, increasing cellular cAMP levels with rolipram mimicked the effects of inhibiting P2RY14, reducing SCP self-renewal *in vitro* and SC proliferation *in* vivo. Our findings in peripheral nerve neurofibromas are entirely consistent with those in NF1 brain tumors. Warrington et al., 2010 found that overexpression *PDE4A1* caused formation of hypercellular lesions with features of NF1-associated glioma, and rolipram inhibited optic glioma growth in an NF1-driven mouse model (Warrington, et al., 2010). Thus, rolipram treatment may be relevant to multiple NF1 manifestations.

In conclusion, purinergic receptor P2RY14 cAMP signaling regulates EGFR-driven Schwann cell precursor self-renewal and nerve tumor initiation in neurofibromatosis. Several groups have suggested that combined inhibition of EGFR and GPCRs could be a promising strategy in cancer therapy (Köse, 2017). P2RY14 is a candidate for this type of therapeutic intervention as a whole-body knockout of this receptor showed no effect on organismal survival or growth (Xu, et al., 2012; Meister, et al., 2014). While the P2RY14 inhibitor (PPTN) has poor PK properties (Robichaud et al., 2011), when drug candidates targeting P2RY14 become available they may be useful for prevention or treatment of NF1-driven neurofibromas.

## Acknowledgements

We thank Takeda Cambridge Limited and Dr. Johannes Grosse for providing the P2RYR14 knock-in mouse. For excellent technical assistance we thank Mark Jackson and Lindsey E. Aschbacher-Smith. J.P-C. was supported by NIH-T32-NS007453 and a Children’s Tumor Foundation Young Investigator Award. This work was supported by grants NIH-R01-NS28840 and NIH-R37-NS083580 to N.R.

## Author Contributions

Conceptualization: N.R., J.P-C. Methodology: J.P-C., J.W., S.K., R.A.C., T.A.R., K.E.C., J.A.C., R.F.H., R.R. Validation: J.P-C., J.W., S.K., R.A.C., R.F.H., S.K. Formal Analysis: J.P-C., W.J., J.A.C., R.R. N.R., Investigation: J.P-C., J.W., S.K. Resources: N.R., R.J.S. Writing-Original Draft: J.P-C., N.R.; Writing-Review & Editing: J.P-C., J W., R.A.C., R.F.H., J.A.C. Visualization: J.P-C., J.W., R.A.F., J.A.C, N.R.; Supervision: N.R. Funding Acquisition: N.R., J.P-C.

## Competing interests

The authors declare no competing interests.

## Methods & Materials

### RESOURCE AVAILABILITY

#### Lead Contact

Further information and requests for resources and reagents should be directed to and will be fulfilled by the Lead Contact, Nancy Ratner, PhD.

#### Materials Availability

This study did not generate new unique reagents.

#### Data and Code Availability Statement

The data sets and original figures generated during this study will be available at Synapse Project (https://www.synapse.org/).

#### Animals

We housed mice in a temperature and humidity-controlled vivarium on a 12hr dark-light cycle with free access to food and water. The animal care and use committee of Cincinnati Children’s Hospital Medical Center approved all animal use. Wild type C57Bl/6 mice were from Jackson Laboratory and were used at 4 and 7 months of age. The *Nf1fl/fl*; *Dhh^+^* mouse line has been described previously by (Wu et al., 2008). We bred *P2RY14-/-* male mice (Meister et. al., 2014; Takeda Cambridge Limited) with *Nf1 fl/fl* female mice to obtain the F1 generation *P2RY14^+/-^;Nf1fl/+*; then, we bred the F1 mice with *DhhCre* male mice to obtain *P2RY14^+/-^;Nf1fl/+*; *Dhh^+^* Then, we bred *P2RY14^+/-^;Nf1fl/+*; *Dhh^+^* (males) with *P2RY14^+/-^;Nf1fl/+*; *Dhh^+^* (females) to obtain *P2RY14^+/-^;Nf1fl/fl*; *Dhh^+^* and *P2RY14^-/-^;Nf1fl/fl*; *Dhh^+^*. We genotyped mice as described by (Meister et al. 2014). Littermates were used for controls. Mice of both sexes were used for all experiments.

#### Human neurofibroma sample collection

Fresh plexiform neurofibromas (n=3) were obtained after medically mandated surgeries. All samples were obtained with patient consent under IRB approval.

#### FACS Analysis

Fresh surgical plexiform neurofibroma (PNF) specimens were enzymatically dissociated as described (Williams et al, 2008). For cell sorting, we incubated cell suspensions with anti-P2RY14 receptor antibody (Rabbit, polyclonal, Alomone labs, # APR-018) on ice for 30 minutes, washed with PBS twice. We then incubated cells with goat-anti-rabbit-APC (Southern Biotech, Cat# 4050-11S), mouse anti-human monoclonal antibodies against p75/NGFR (Becton-Dickinson, Cat# 40-1457) bound to phycoerythrin (PE), and EGFR (Fitzgerald, Acton, MA, Cat# 61R-E109BAF,) bound to FITC on ice in PBS/ 0.2% human serum for 30 min. After washing, we re-suspended cells in PBS/ 0.2% human serum containing 2 µg/mL 7-aminoactinomycin D (7-AAD) (Invitrogen, Cat# A1310). We carried out isotopic controls with irrelevant mouse IgG1-APC, mouse-IgG1-PE and mouse-IgG1-FITC in parallel. Cells were FACS-sorted using a four-laser FACSDiva (Becton-Dickinson) to acquire alive Schwann cell sub-population (P75^+^/EGFR^+^/P2RY14^+^/7-AAD^-^ and P75^+^/EGFR^+^/P2RY14^-^/7-AAD^-^). Three primary human PNFs were FACS-sorted independently.

#### Western blotting

Primary antibodies and dilutions were: Anti-Purinergic Receptor P2RY14 rabbit (extracellular) (1:200, Sigma Aldrich, Cat#: P0119); GPR105 Polyclonal antibody rabbit (1:200, Thermo Fisher, Cat#: PA5-34087); phospho PKA substrate (RRXS*/T*) (100G7E) rabbit (1:1000, Cell Signaling Technology, Cat#: 9624S); phosphor-p44/42 MAPK (Erk1/2) (Thr202/Tyr204) rabbit (1:1000, Cell Signaling Technologies, Cat#: 8544S); P44/42 MAPK (Erk1/2) (137F5) rabbit mAB (1:1000, Cell Signaling Technologies, Cat#: 4695S).

#### Sphere culture

We dissociated DRG from E12.5 embryos with 0.25% Trypsin (Thermo Fisher, Cat# 25200056) for 20 min at 37 ÅãC and obtained single-cell suspensions with narrow-bore pipettes and a 40 μm strainer (BD-Falcon), plating the cells in 24 well low attachment plates (Corning). The free-floating cells were cultured in serum-free medium with EGF and FGF as described (Williams et al. 2008). For passage, we dissociated spheres with 0.05% Trypsin (Thermo Fisher, Cat# 25300054) at 37Åã for 5 min. For shRNA treatment and sphere counts, we plated SCP cells at low density to avoid sphere fusion (1000 cells/well in 24 well plates). In the P2RY14 drug studies, we treated the cells with P2RY14 inhibitor: P2RY14 Antagonist Prodrug 7j hydrochloride (Axon Medchem, Cat# 1958) at concentrations of 30nM, 100nM, 300nM and 500nM. Three biological replicates produced similar results. Dose response analysis confirmed that the optimal concentration of PPTN was 300nM in this assay. For the shRNA experiments, we treated cells with three different lentiviral particles: shRNA control plasmid DNA (Sigma-Aldrich, Cat# SHC016-1EA), shP2RY14(09) (Sigma-Aldrich, Cat# SHCLNG-NM_133200; TRCN0000328609), shP2RY14(84) (Sigma-Aldrich, Cat# SHCLNG-NM_133200; TRCN0000328684), shP2RY14(64) (Sigma-Aldrich, Cat# SHCLNG-NM_133200; TRCN0000026664) at MOI = 10, 24 h after plating. For rolipram experiment, we treated cells with 1uM rolipram (Selleck, Cat# S1430). To passage, we centrifuged sphere cultures, treated with 0.05% Trypsin for 3 min., dissociated and plated at 2 × 10^4^ cells/ml in 50% conditioned and 50% fresh medium. We counted secondary spheres after 14 days. For every cell line three biological replicates with three technical replicates were done. Of the three biological replicates the best one was reported as a representative (n=3). Spheres were counted with an inverted phase contrast microscope after 6 days of plating.

### Primary SC culture

Embryonic day 12.5 primary mouse SCs (mSC) were isolated from dorsal root ganglia (DRG) with neuronal contact in N2 medium with nerve growth factor, then removed from neurons and cultured in SC media (DMEM (Thermo Fisher, Cat# 11965118) + 10% FBS (Gemini Bio-Products, Cat# A87G02J) + β-heregulin peptide (R&D Systems, Cat# 396-HB-050) + forskolin (Cayman Chemical, Cat# 11018)) for 1–3 passages as described (Kim, Roseanbaum, Marchionni, Ratner, & DeClue, 1995).

### CRISPR/Cas9 immortalized human SCs (IHSC-1λ)

The immortalized human Schwann iHSC-1λ were infected with lentivirus derived from pLentiCRISPRv2 puro (Addgene, Cat#78852) expressing Cas9 and either a scrambled gRNA or one directed against the human NF1 gene designed to cleave between amino acids 157 and 158 in Exon 4. Pooled puromycin resistant cells were evaluated for NF1 expression by western blot (Bethyl Laboratories, Cat#A300-140A) and growth and EdU incorporation assays were performed. gRNA #1: 5’-CAGTCTTTAGTCGCATTTCTACC-3’: Target Exon 4, cut site between aa 157 and 158. gRNA #2: 5’-ACACTGGAAAAATGTCTTGC-3’: Target Exon 3, cut site at aa 94.

### Immortalized SCs P2RY14 mechanistic experiment

Immortalized human SCs were treated plated in 6-well plates (200,000 cells per well) in DMEM +10%FBS + P/S (Penicillin). After cells were confluents, cells were treated with serum free media for 4 hours. At the 3-hour time-point, cells were treated with 200uM IBMX (Millipore Sigma, Cat.# I5879) for 1 hour. At the 3.5-hour time-point, cells were treated with the 500nM PPTN inhibitor for 30 minutes. At the 4-hour time-point, cells were stimulated with 10uM UDP analog MRS2690 (Tocris, Cat. #: 2915) for 15 mins. Cells were collected in RIPA buffer, denatured and used to run western blot. For every cell line three biological replicates with three technical replicates were done. Of the three biological replicates the best one was reported as a representative (n=3).

### Direct cAMP ELISA assay

Primary Schwann cells were plated on poly-L-lysine coated 6 well plate (∼750,000 cells per well) in DMEM + 10 % FBS + β HRG + forskolin. After cells were confluent, cells were starved with serum-free N2 media with N2 supplement and left incubating for 16 hours. Cells were pre-incubated with MEK inhibitor for two hours prior to stimulation. Stimulation was carried out for 2 minutes and cAMP measurement was done according to manufacturer’s protocol (Direct cAMP ELISA kit by Enzo Life Sciences, Cat# ADI-900-066) using the acetylated format. An aliquot prior to cAMP measurements was set aside for protein quantification using the Bio-Rad protein assay kit. For every cell line two biological replicates with three technical replicates were done. Of the two biological replicates the best one was reported as a representative (n=3 biological replicates, 3 technical replicates).

### Immunostaining

For frozen sections, OCT was removed by incubation with 1XPBS. We permeabilized cells in ice cold MeOH for 10 min., followed by incubation in normal donkey serum (Jackson ImmunoResearch Cat# 017–000-121) and 0.3% Triton-X100 (Sigma-Aldrich Cat# X100). Ki67 (1:200, Cell Signaling Technologies Cat# 12202S), anti-CNPAse (1:250, Sigma Aldrich Millipore, Cat# AB9342); β-galactosidase polyclonal antibody (1:1000, Thermo Fisher, Cat#: A-11132); β-actin (13E5) Rabbit mAb (HRP Conjugate) (1:1000, Cell Signaling, Cat#: 5125S). All secondary antibodies were donkey anti Rat/Rabbit/Goat from Jackson ImmunoResearch, reconstituted in 50% glycerol and used at 1:250 dilution. To visualize nuclei, sections were stained with DAPI for 10min., washed with PBS and mounted in FluoromountG (Electron Microscopy Sciences, Hatfield, PA). Images were acquired with ImageJ Acquisition software using a fluorescence microscope (Axiovert 200M) with 10x/0.4 or 40x/0.6 objectives (Carl Zeiss, Inc.), or with NIS-Elements software using confocal microscopy (Nikon).

### Electron Microscopy

Mice were perfusion fixed with 4% paraformaldehyde and 2.5% glutaraldehyde in 0.1-M phosphate buffer at 7.4 pH. Saphenous nerve was dissected out and postfixed overnight, then transferred to 0.175 mol/L cacodylate buffer, osmicated, dehydrated, and embedded in Embed 812 (Ladd Research Industries). Ultrathin sections were stained in uranyl acetate and lead citrate and viewed on a Hitachi H-7600 microscope.

### Mouse dissection and quantification of neurofibroma number and size

To quantify neurofibroma number and size, we perfused mice and used a Leica dissecting microscope to dissect the spinal cord with attached DRG and nerve roots at the ages of 4-months and 7-month, as previously described (Wu, et al., 2016). A neurofibroma was defined as a mass surrounding the DRG or nerve roots, with a diameter greater than 1mm, measured perpendicular to DRG/nerve roots. Neurofibroma diameter for each mouse were measured with Image J.

### RT-PCR

We isolated total RNA from WT and Nf1-/-spheres treated with shP2RY14 using the RNeasy Plus Micro-Kit (QIAGEN, Cat# 74034) and made cDNA using the High Capacity Reverse Transcription Kit (Thermo Fisher, Cat# 4368813). We conducted rt-PCR as described in (Williams et al. 2008). Primer sequence: P2RY14 forward: 5’-AGCAGATCATTCCCGTGTTGT-3’. P2RY14 reverse: 5’-TCTCAAGAACATAGTGGTGGCT-3’.

### Genotyping

Power SYBR Green PCR Master Mix (Thermo Fisher, Cat# 4368702) was used for genotyping. We genotyped mice as described by (Meister et al. 2014). Genotyping primers: b-galactosidase sense: 5’-AGAAGGCACATGGCTGAATATCGA-3’. P2RY14 forward: (5’-AGCTGCCGGACGAAGGAGACCCTGCTC-3’. P2RY14 reverse: 5’-GGTTTTGGAAACCTCTAGGTCATTCTG-3’ (Meister et al., 2014).

## Statistics

Statistical parameters, including the type of tests, number of samples (n), descriptive statistics and significance are reported in the figures and figure legends. Two-group comparisons used Student’s t-tests. When single agents were tested at different concentration in a single cell type, we used a one-way ANOVA with a Dunnett’s multiple comparisons test. When multiple genotypes were analyzed in a single experiment, we used a Two-way ANOVA with multiple comparisons, without matching, and correction with the Holm-Sidak test. Mann-Whitney test was used for comparisons between genotypes for tissue widths and neurofibroma incidence (GraphPad Prism V9). All data unless otherwise stated is represented as average ± SD, and was analyzed in GraphPad Prism 7.

---

**Supplemental figure 1:**
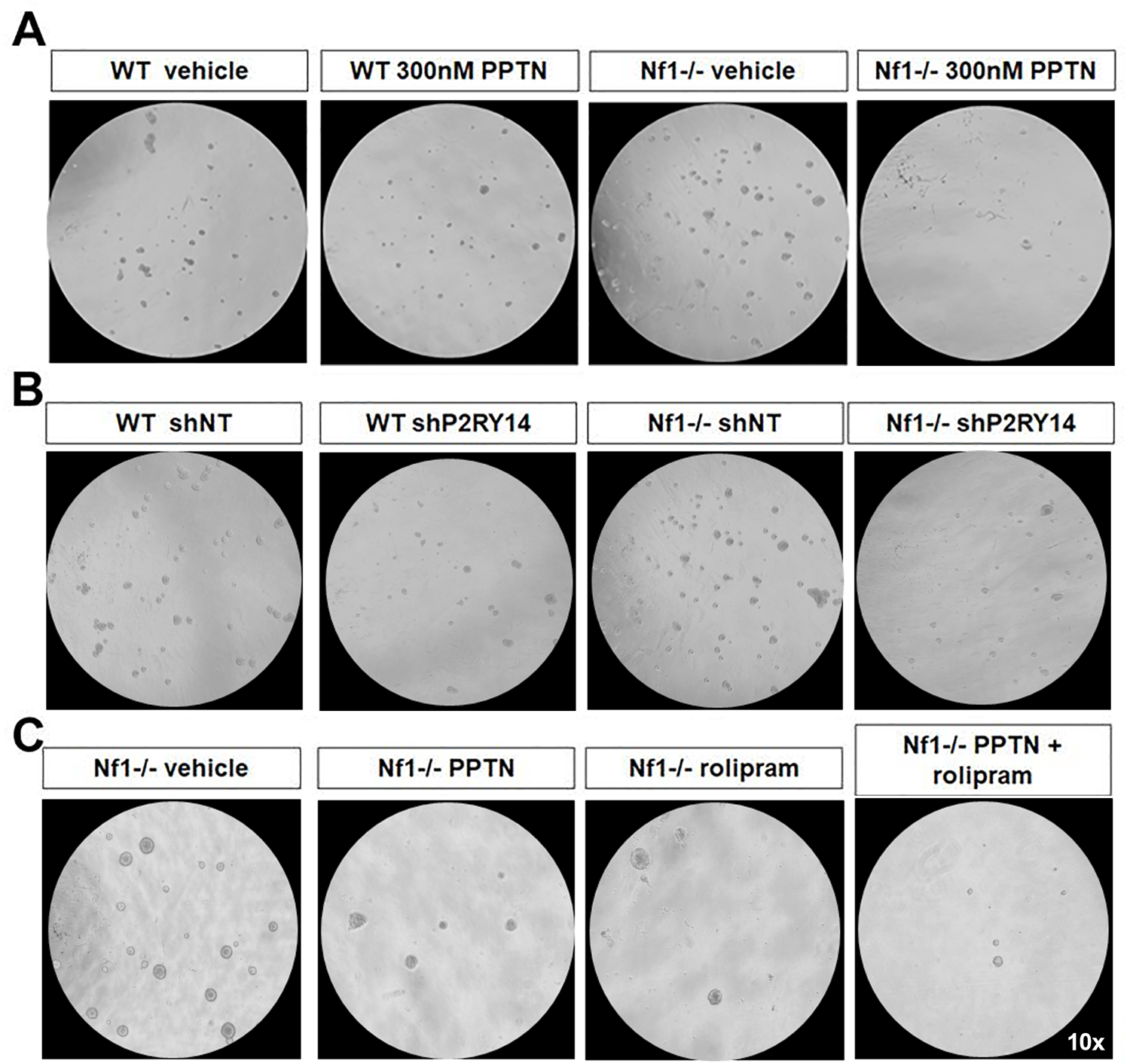
Photomicrographs of three different experiments in which *WT* and *Nf1-/-* SCPs were treated with PPTN, shP2RY14 and rolipram. A) Photomicrographs of *WT* and *Nf1-/-* mouse SC spheres treated with the selective P2RY14 inhibitor (PPTN; 300nM). B) Photomicrographs of *WT* and *Nf1-/-* mouse SC spheres treated with shNT and shP2RY14 (09). C) Photomicrographs of *NF1-/-* SCPs treated with vehicle and 1uM of rolipram.

**Supplemental figure 2:**
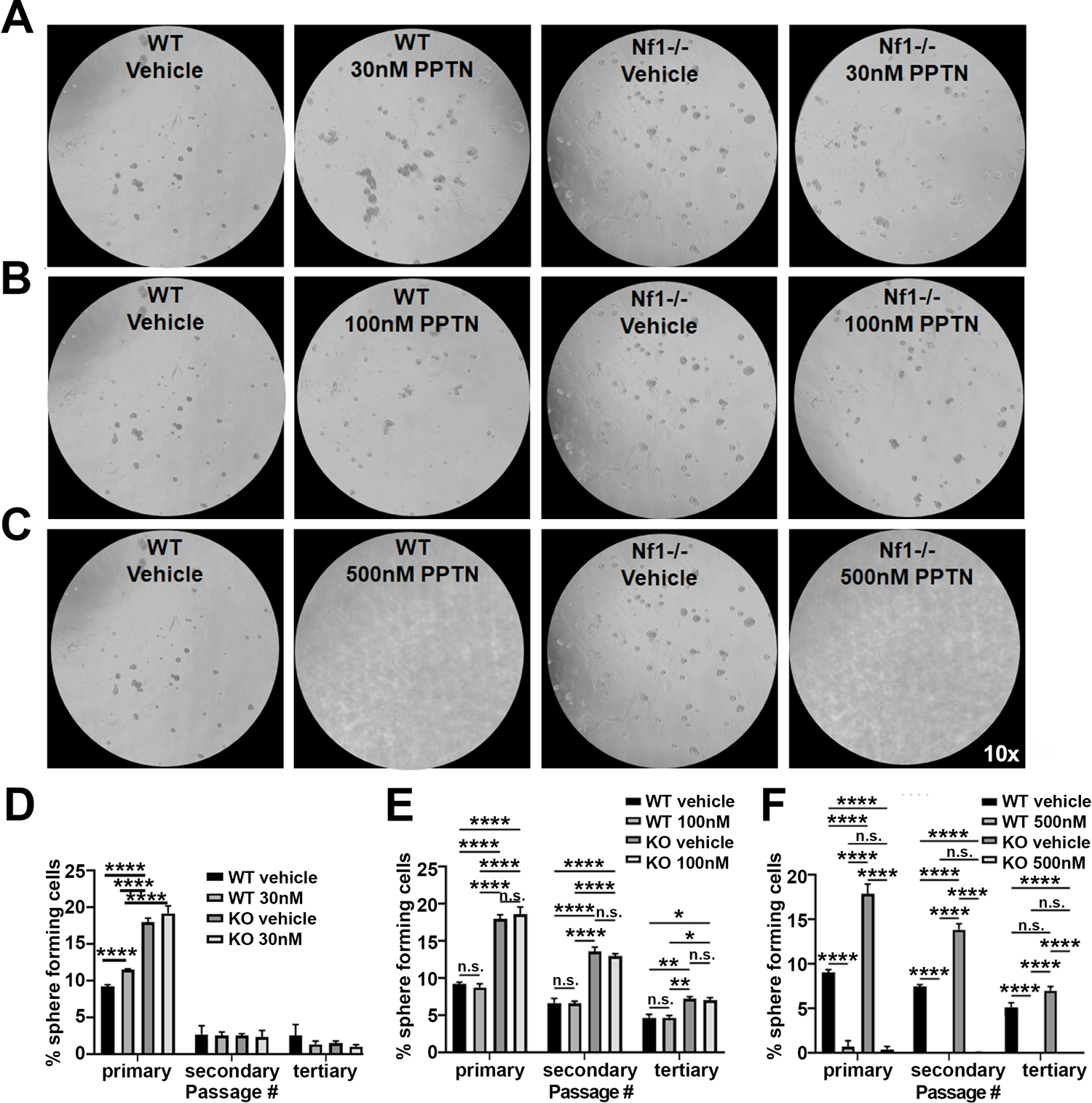
PPTN dose response analysis in *WT* and *Nf1-/-* SCPs. A) Photomicrographs of *WT* and *Nf1-/-* mouse SC spheres treated with the P2RY14 inhibitor (PPTN; 30nM). B) Photomicrographs of *WT* and *Nf1-/-* mouse SC spheres treated with the P2RY14 inhibitor (PPTN; 100nM). C) Photomicrographs of *WT* and *Nf1-/-* mouse SC spheres treated with the P2RY14 inhibitor (PPTN; 500nM). (Note: for figures S2A-C, the same WT and *Nf1-/-* vehicle controls photomicrographs are used for each dose). D) Quantification of the percent of sphere forming cells after treatment with the P2RY14 inhibitor (PPTN; 30nM) in *WT* and *Nf1-/-* mouse SC spheres (two-way ANOVA: primary and secondary passage: ****p<0.0001, tertiary passage: *p=0.0168, **p=0.0074) E) Quantification of percent sphere forming cells after treatment with the P2RY14 inhibitor (PPTN;100nM) in *WT* and *Nf1-/-* mouse SCP (two way ANOVA: primary, secondary and tertiary passage: ****p<0.0001). F) Quantification of percent sphere forming cells after treatment with the P2RY14 inhibitor (PPTN; 500nM) in *WT* and *Nf1-/-* mouse SCP; results show that this concentration was toxic to the cells (two way ANOVA: primary, secondary and tertiary passage: ****p<0.0001).

**Supplemental figure 3:**
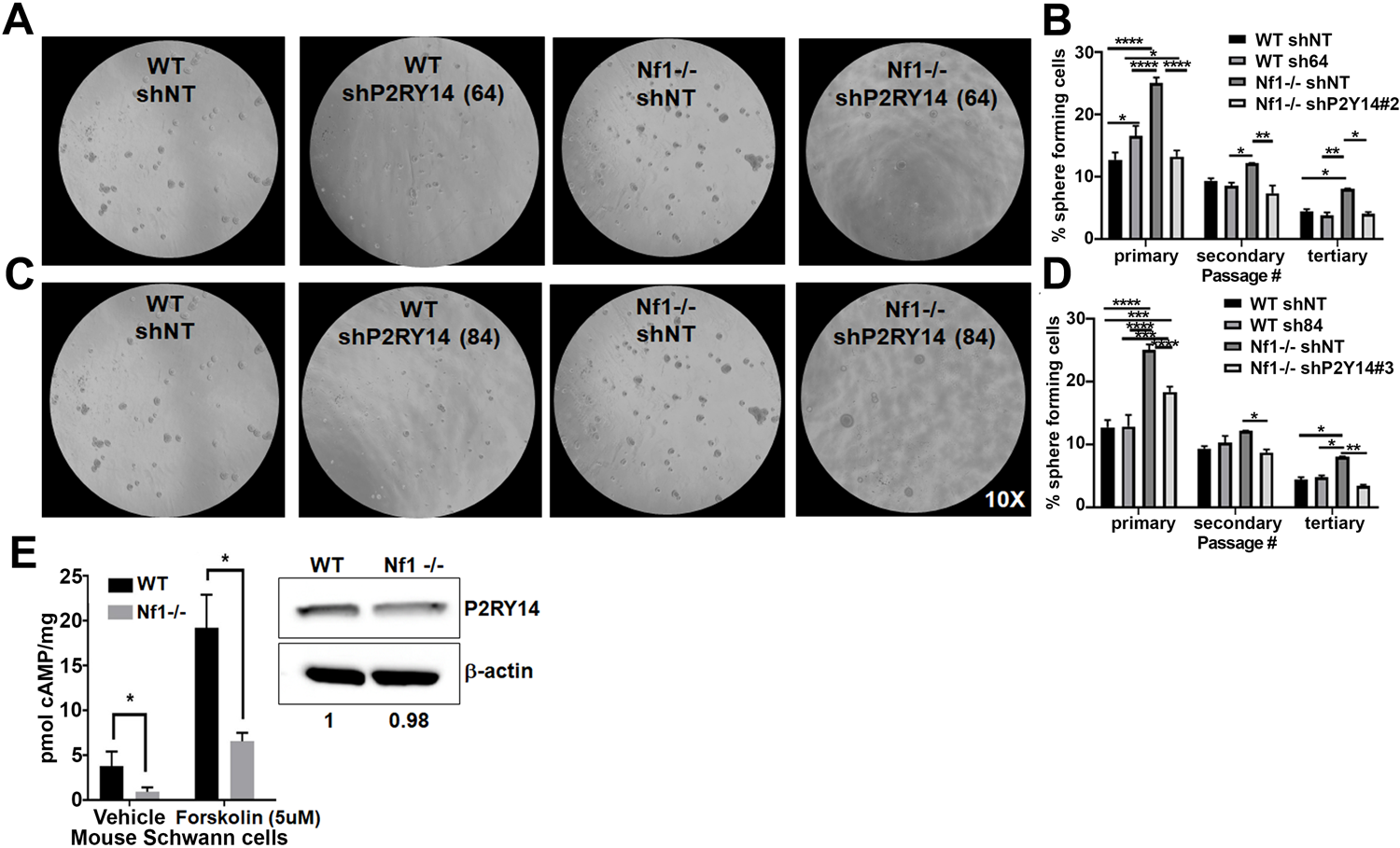
Photomicrographs and quantification analysis of two biological replicates of shP2RY14 in *WT* and *Nf1-/-* SCPs. A) Photomicrographs of *WT* and *Nf1-/-* SCP treated with shP2RY14 (64). B) Quantification of the percent of sphere forming cells in *WT* and *Nf1-/-* SCP after treatment with shP2RY14 (64) (n=3; two-way ANOVA: primary: *p=0.0184, ****p<0.0001; secondary: *p=0.0270, **p=0.0024; tertiary: *p=0.0270, **p=0.0078). C) Photomicrographs of *WT* and *Nf1-/-* SCP treated with shP2RY14 (84). D) Quantification of the percent of sphere forming cells in *WT* and *Nf1-/-* SCP after treatment with shP2RY14 (84) (n=3; two-way ANOVA: primary: ***p=0.0005, ****p<0.0001; secondary: *p=0.0358; tertiary: *p=0.0245, **p=0.0030). (Note: for figures S3A & S3C, the same WT and *Nf1-/-* shNT control photomicrographs are used). E) *WT* and *Nf1-/-* E12.5 mouse SCs cAMP levels measured at baseline and after activation of AC via forskolin stimulation (n=3, multiple t-test: vehicle *p=0.0407, forskolin (5uM) *p=0.0045).

**Supplemental figure 4:**
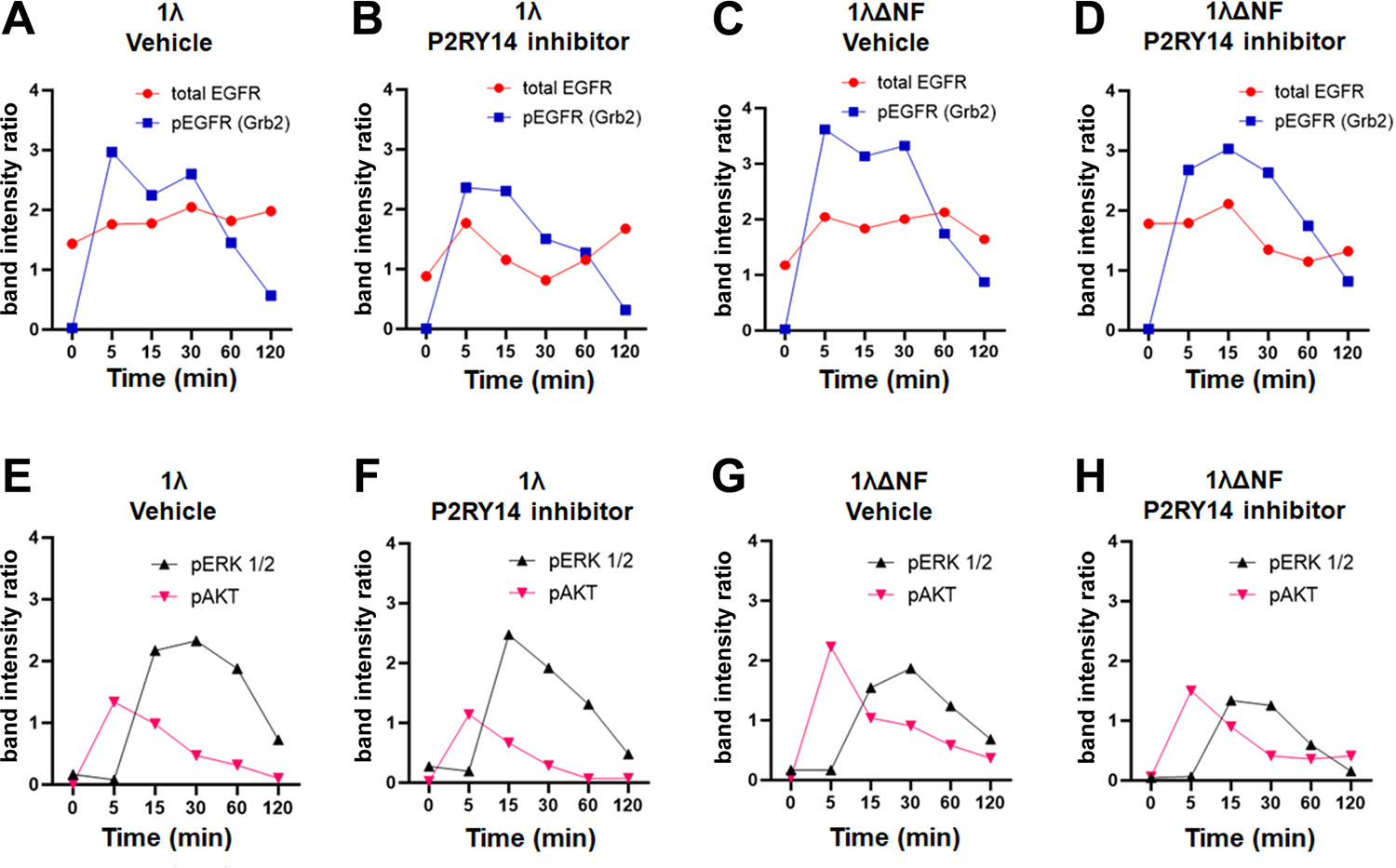
Quantification of 1λ and 1λΔNF1 cells treated with PPTN and EGF. A) Graph shows total EGFR and pEGFR (Grb2) dynamics overtime after addition of EGF in vehicle treated 1λ cells. B) Graph shows total EGFR and pEGFR (Grb2) dynamics overtime after addition of EGF in P2RY14 inhibitor (PPTN) treated 1λ cells. C) Graph shows total EGFR and pEGFR (Grb2) dynamics overtime after addition of EGF in vehicle treated 1λΔNF1 cells. D) Graph shows total EGFR and pEGFR (Grb2) dynamics overtime after addition of EGF in P2RY14 inhibitor (PPTN) treated 1λΔNF1 cells. E) Graph shows pERK1/2 and pAKT dynamics overtime after addition of EGF in vehicle treated 1λ cells. F) Graph shows pERK1/2 and pAKT dynamics overtime after addition of EGF in P2RY14 inhibitor (PPTN) treated 1λ cells. G) Graph shows pERK1/2 and pAKT dynamics overtime after addition of EGF in vehicle treated 1λΔNF1 cells. H) Graph shows pERK1/2 and pAKT dynamics overtime after addition of EGF in P2RY14 inhibitor (PPTN) treated 1λΔNF1 cells.

**Supplemental figure 5:**
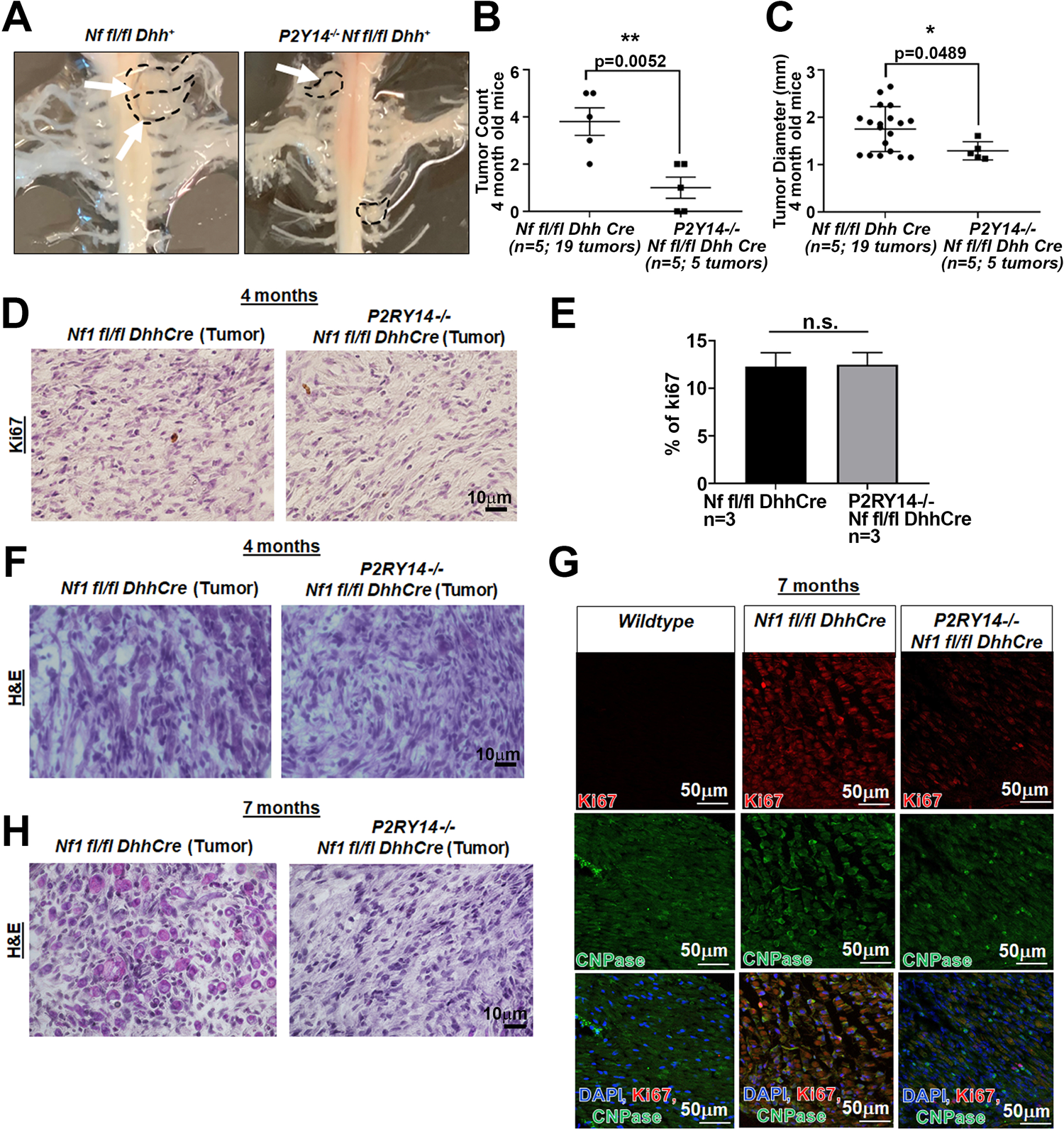
Tumor dissection of *Nf1fl/fl*;*Dhh+* and *P2RY14^-/-^;Nf1fl/fl*;*Dhh+* mice at 4-months and immunostaining analysis of 4-months and 7-months sciatic nerve and tumors. A) Representative image of gross dissection of *Nf1fl/fl*;*Dhh+* and *P2RY14^-/-^;Nf1fl/fl*;*Dhh+* mice at 4-months. B) Neurofibroma tumor number quantification at 4-months of age (unpaired t-test **p=0.0052). C) Neurofibroma tumor diameter quantification at 4-months (unpaired t-test **p=0.0489) (for figures B & C: *Nf1fl/fl*;*Dhh+* n=5 mice, 19 neurofibroma tumors; *P2RY14-/-;Nf1fl/fl*;*Dhh+* n=5 mice; 5 neurofibroma tumors). D) Ki67 staining of *Nf1fl/fl*;*Dhh+* and *P2RY14-/-;Nf1fl/fl*; *Dhh+* neurofibromas at 4-months of age. E) Quantification of percent of Ki67 positive cells in *Nf1fl/fl*;*Dhh+* and *P2RY14-/-;Nf1fl/fl*;*Dhh+* 4-months old mice (t-test; n.s.). F) H&E staining at 4-months in *Nf1fl/fl*;*Dhh+* and *P2RY14-/-;Nf1fl/fl*;*Dhh+* mice. G) Ki67 immunofluorescence co-labeling with CNPase (SC marker) in 7-months sciatic nerve. H) H&E tumor staining at 7-months in *Nf1fl/fl*;*Dhh+* and *P2RY14-/-;Nf1fl/fl*;*Dhh+* mice.

**Supplemental figure 6:**
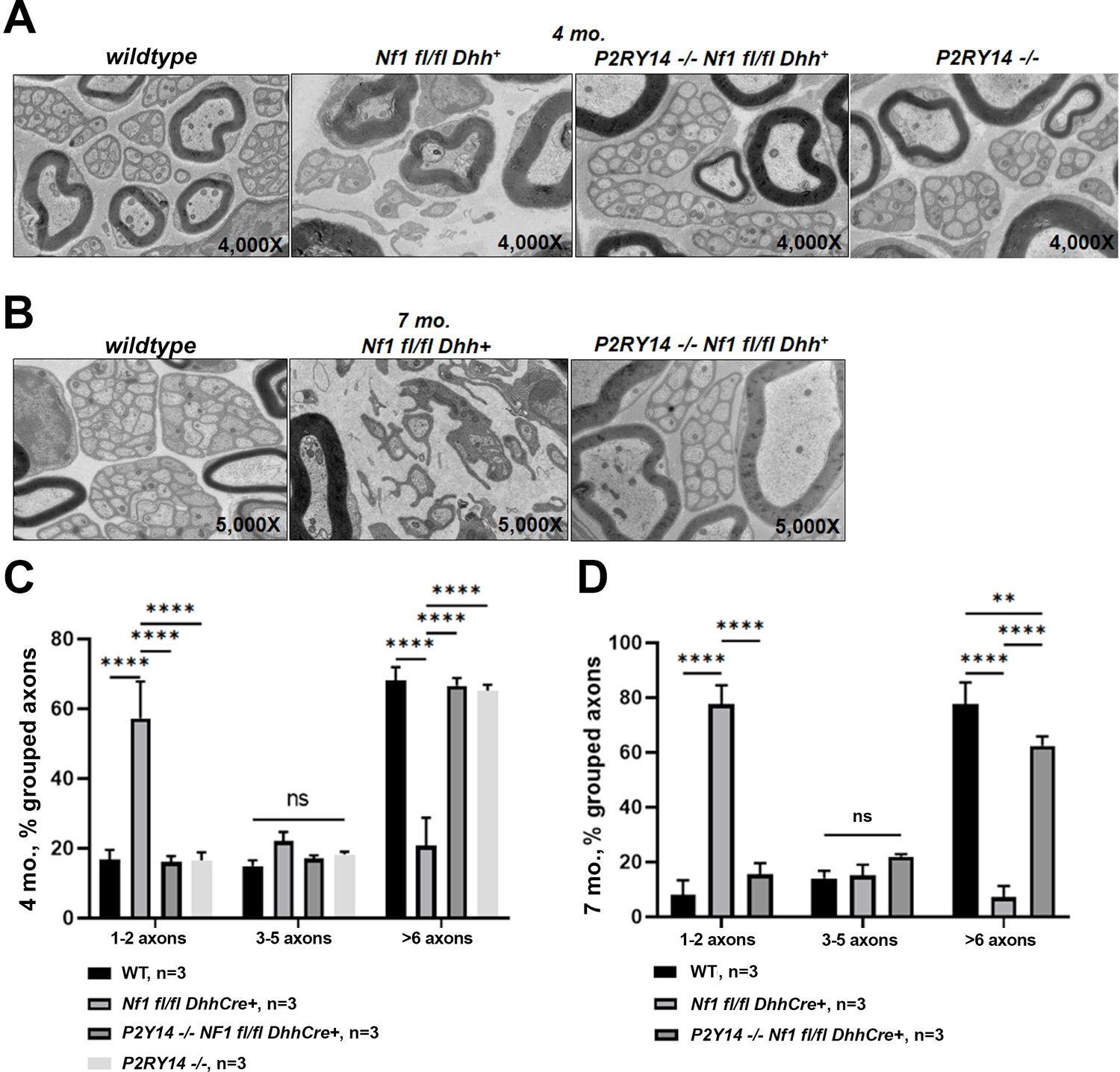
P2RY14 deletion improves nerve ultrastructure. A) Electron micrograph of saphenous nerve of 4-months *WT, Nf1fl/fl*;*Dhh, P2RY14^-/-^;Nf1fl/fl*;*Dhh+* and *P2RY14^-/-^* mice. B) Electron micrograph of saphenous nerve of 7-months *WT, Nf1fl/fl*;*Dhh* and *P2RY14^-/-^;Nf1fl/fl*; *Dhh+* mice. C) Remak bundle quantification at 4-months of age (n=3; two way ANOVA: ****p<0.0001). D) Remak bundle quantification at 7-months of age (n=3; two way ANOVA: **p=0.0027, ****p<0.0001).

## Notes

### Competing Interest Statement

The authors have declared no competing interest.

